# Coordinated Immune Cell Networks in the Bone Marrow Microenvironment Define the Graft versus Leukemia Response with Adoptive Cellular Therapy

**DOI:** 10.1101/2024.02.09.579677

**Authors:** Katie Maurer, Cameron Y. Park, Shouvik Mani, Mehdi Borji, Livius Penter, Yinuo Jin, Jia Yi Zhang, Crystal Shin, James R. Brenner, Jackson Southard, Sachi Krishna, Wesley Lu, Haoxiang Lyu, Domenic Abbondanza, Chanell Mangum, Lars Rønn Olsen, Donna S. Neuberg, Pavan Bachireddy, Samouil L. Farhi, Shuqiang Li, Kenneth J. Livak, Jerome Ritz, Robert J. Soiffer, Catherine J. Wu, Elham Azizi

**Author notes:** These authors contributed equally (Katie Maurer, Cameron Y. Park). Senior authors (Catherine J. Wu, Elham Azizi).

## Abstract

Understanding how intra-tumoral immune populations coordinate to generate anti-tumor responses following therapy can guide precise treatment prioritization. We performed systematic dissection of an established adoptive cellular therapy, donor lymphocyte infusion (DLI), by analyzing 348,905 single-cell transcriptomes from 74 longitudinal bone-marrow samples of 25 patients with relapsed myeloid leukemia; a subset was evaluated by protein-based spatial analysis. In acute myelogenous leukemia (AML) responders, diverse immune cell types within the bone-marrow microenvironment (BME) were predicted to interact with a clonally expanded population of *ZNF683^+^GZMB^+^* CD8+ cytotoxic T lymphocytes (CTLs) which demonstrated *in vitro* specificity for autologous leukemia. This population, originating predominantly from the DLI product, expanded concurrently with NK and B cells. AML nonresponder BME revealed a paucity of crosstalk and elevated *TIGIT* expression in CD8+ CTLs. Our study highlights recipient BME differences as a key determinant of effective anti-leukemia response and opens new opportunities to modulate cell-based leukemia-directed therapy.

## Introduction

Despite the clinical successes of cancer immunotherapies such as immune checkpoint blockade (ICB) and adoptive cellular therapies, we lack a clear understanding of the specific role and characteristics of tumor-infiltrating immune cells in the local microenvironment.^1–5^ Coordinated response among various cellular players in the tumor microenvironment (TME) may be a key component of effective tumor-directed responses.^6–13^ Responsiveness of patients to diverse immunotherapeutic modalities varies not only from malignancy to malignancy, but also across patients within an individual cancer type^14–16^ — underscoring the need to define the molecular and cellular determinants of response in a disease-specific context.

The graft-versus-leukemia (GvL) effect is a key feature of maintenance of remission following allogeneic hematopoietic stem cell transplantation (HSCT), wherein donor immune cells eliminate residual malignant cells in the recipient. Disease relapse after HSCT represents immunologic escape by malignant cells, after which GvL can be reinvigorated by infusion of donor lymphocytes (DLI) from the original stem cell donor.^17–19^ Despite long-standing clinical recognition of this phenomenon, the mechanisms of GvL remain largely undefined. Further, response to DLI is highly variable among hematologic malignancies, with 70-80% of patients with chronic myelogenous leukemia (CML) demonstrating response to DLI, contrasted to only 15-20% of patients with acute myelogenous leukemia (AML) who respond.^20^ In this context, DLI offers the opportunity to carefully examine the cellular determinants of response and resistance to GvL in the leukemia bone marrow microenvironment (BME).^21–25^

Applications of single-cell resolution transcriptomic and spatial profiling are transforming our ability to disentangle the complexity of the TME in patient specimens.^26–32^ Multi-omic sequential profiling of patient specimens and computational frameworks now allow integration of heterogeneous high-dimensional datasets across patients and time points before and after therapy while accounting for temporal dependencies and confounding factors inherent to clinical data^4,31^. Indeed, we previously applied single cell transcriptome (scRNA-seq) analysis and novel machine learning methods to study the temporal dynamics of marrow-infiltrating T-cell subpopulations collected from CML patients treated with DLI.^5,33^ In that study, we identified the association between expansion of precursor exhausted T cells (T_PE_) and favorable clinical outcome, while a subset of terminal exhausted T cells (T_EX_) was enriched prior to DLI only in responders^5^. These observations now motivate not only a more comprehensive characterization of the entire leukemic marrow microenvironment to more broadly investigate crosstalk between heterogeneous leukemic states and immune cells in shaping the GvL response, but also a focus on the high-priority and challenging setting of relapsed AML for which few therapies exist and outcomes are poor.^34^

Herein, we present an integrated experimental and computational framework for unbiased characterization of spatiotemporal dynamics of cell states and complex intercellular interactions in the BME of leukemia patients. Using DLI for relapsed leukemia as an informative model system of immunotherapy, we define cellular drivers of GvL response versus resistance by single-cell transcriptomic, proteomic (Cellular indexing of transcriptomes and epitopes - CITE) and T cell receptor (TCR) profiling of 74 bone marrow specimens collected longitudinally from 9 AML and 16 CML patients. We developed DIISCO (Dynamic Intercellular Interactions in Single Cell transcriptOmics)^35^, a machine learning framework for time-resolved analysis of cell-cell interactions that demonstrated that CD8+ CTL anti-leukemia activity is closely linked to dynamics of other immune cell states. We confirmed these findings through spatial mapping of cells in the context of the marrow microenvironment in the same AML patients and with functional *in vitro* studies in 6 AML responders. Altogether, our findings indicate the requirement for coordinated multicellular interactions for response to adoptive cellular immunotherapy for myeloid leukemia. Our machine learning-enabled spatiotemporal modeling to pinpoint key immune networks in clinical specimens, followed by *in vitro* and large cohort validation to confirm their biomarkers, provides a roadmap for broader studies of immunotherapy response and resistance.

## Results

### Distinct marrow microenvironments among myeloid leukemias

To dissect the composition of immune cell states in the BME of leukemia patients, we profiled the transcriptomes of individual marrow cells originating from 5 responders (R) and 4 nonresponders (NR) to DLI administered for treatment of post-HSCT relapsed AML (**Table 1**). Patients in the R and NR groups had similar baseline characteristics including HLA-match of donor, recipient/donor sex, time from HSCT to relapse, time from HSCT to DLI, and timing of pre-DLI samples (**Figure S1A,B**; **Table 1**). R patients had a longer duration of post-DLI marrow sampling due to longer survival after DLI. Patients in both groups had comparable disease burden at relapse, defined as percent of bone marrow blasts (**Figure S1C**). Three of five Rs and two of four NRs received cytoreducing therapy between relapse and DLI in an attempt to achieve remission prior to DLI (**Table 1**). Two Rs and one NR had minimal disease (i.e. <5% blasts but marrow dysplasia and/or progressive cytopenias and/or recurrence of mutations by next-generation sequencing) at relapse and did not receive therapy prior to DLI given the minimal disease burden. All patients except one NR had <5% blasts at the time of DLI. The exception to this was NR 3512 whose disease at relapse was considered as smoldering and therefore DLI was pursued without prior therapy despite 16% marrow blasts.

All patients received between 1-3 doses of DLI (median 1 for R and 1.5 for NR) with a median dose of 1×10^7^ CD3+ cells/kg for R vs 1.6×10^7^ CD3+ cells/kg for NR (**Table 1**). Response was defined as durable morphologic and molecular remission for at least one year after DLI, while NRs lacked post-treatment reduction in disease burden. A median of 3 (range 2-6) serial marrow biopsy specimens collected before and after DLI were evaluated per patient (**Figure 1A**; **Table S1)**. A total of 47,372 viable mononuclear cells originating from 32 BM samples from the 9 subjects were transcriptionally profiled by scRNA-seq and scTCR-seq, and by scCITE-seq for proteomic characterization. In addition, we performed these same characterizations on DLI product samples of 4 AML-R (patients 3501, 3503, 3505, and 3507) and 3 AML-NR (patients 3512, 3512, 3516) from the discovery cohort, yielding 43,918 high-quality cells for analysis, including 21,936 T-cells (**Figure 1B**). As controls, we similarly profiled 7,617 marrow mononuclear cells collected from two post-HSCT AML patients who did not relapse, as well as 27,131 marrow mononuclear cells collected pre- and post-treatment from three AML patients who relapsed following HSCT and entered remission following chemotherapy but did not receive DLI (denoted chemotherapy only/no DLI controls).

**Figure 1.**
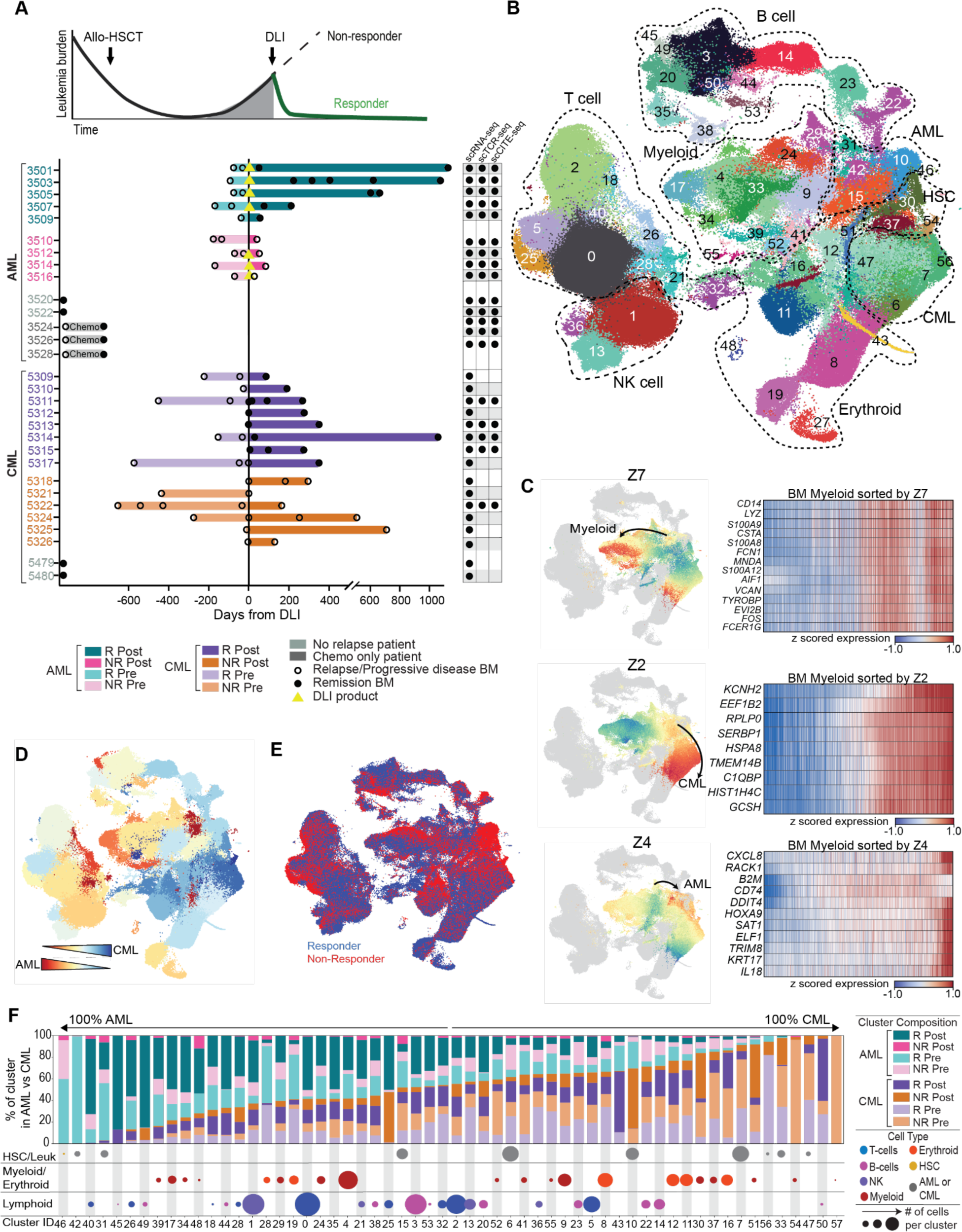
Experimental design and global map of leukemic bone marrow microenvironment. **A.** Schema of marrow samples analyzed by scRNA-seq, collected from patients with AML or CML, before or after DLI. Leukemia burden (y axis) was quantified by bone marrow blasts (AML) or *BCR-ABL* transcript (CML). All AML samples and a subset of CML samples were also analyzed by scTCR-seq and scCITE-seq (right). Vertical center line - day 0 of first DLI infusion. Pre-DLI samples (lighter bars) and post-DLI samples (darker bars) were analyzed for both responders (AML: teal, CML: purple) and nonresponders (AML: pink, CML orange). Two control patients who did not relapse were included for both AML and CML (gray) as well as three AML patients with post-HSCT relapse who entered remission after chemotherapy without DLI (sage). Available DLI infusion products were also analyzed by scRNA, scTCR, and scCITE-seq (yellow triangles). **B.** 2D UMAP projection of all 461,940 transcriptomes from AML, CML, and control samples. Each dot represents a cell and is colored based on 58 clusters denoting distinct cell subsets. Clusters 32, 34, 35, 43, and 53, were removed from analysis as doublets, resulting in 53 distinct cell subsets and 451,533 cells. Dashed lines encompass major cell types. **C.** Coloring of the 2D UMAP for BM myeloid cells based on Decipher latent factors (**Methods**), which reveal distinct trajectories for healthy myeloid differentiation (Z7, top), CML evolution (Z2, middle), and AML evolution (Z4, bottom). Grey - non-myeloid cells. Heatmaps on right - gene expression profiles of BM myeloid cells sorted by each Decipher component per trajectory. **D.** Coloring of the clusters in the 2D UMAP based on percent of cells per cluster from each disease/patient cohort. Red: Clusters composed of only cells from AML patients; blue: clusters composed of only cells from CML patients. **E.** Coloring of the 2D UMAP based on patient response outcome. **F.** *Top:* Bar graph normalized by total number of cells per disease type demonstrating the percent of each cluster from AML responders (teal), AML nonresponders (pink), CML responders (purple), and CML nonresponders (orange), as well as percent of cluster from pre-DLI (lighter colors) or post-DLI (darker colors) time points. *Bottom:* Relative size of clusters (increasing size of dots indicates more cells in cluster) and major cell type per cluster, including HSC/leukemia cells (top row, HSC: gold, leukemia cells: gray), myeloid/erythroid cells (middle row, myeloid: red, erythroid: orange); and lymphoid cells (bottom row, T cells: blue, B cells: dark pink, NK cells: purple).

To expand our compendium of marrow-derived mononuclear cells from patients with myeloid malignancies, we merged the AML marrow data with our previously generated dataset from 42 marrow specimens from 16 DLI-treated patients with CML^5^ and extended the analysis from T cells to the entire leukemic marrow microenvironment. We constructed an integrated atlas from all marrow and mononuclear cells (using scVI^36,37^ batch correction, see **Methods**) from these 25 patients for a total of 451,553 high-quality transcriptomes from individual immune cells (mean of 4,560 cells per sample; **Table S2**; **Figure S1D,E**). Using Phenograph clustering^38^ on scVI latent components and removing doublets, we identified 53 distinct cell states, including subsets of T, B, NK, monocyte, hematopoietic progenitor, and leukemia cells (**Figure 1B**; **Methods**). Forty-three cell states (81%) were not predominated by a specific sample, i.e. had <60% cells from one sample (**Methods**; **Figure S1E**) and replicates from the same specimen profiled with different sequencing chemistries exhibited substantial overlap (**Figure S1D**), confirming successful correction of technical variation, while preserving patient differences. Of the 10 (19%) clusters dominated by a single sample, 5 were leukemia/HSC-enriched as expected (C10, C42, C46, C54, C56), and 5 contained immune cells (i.e. C18 - CD4 T progenitor exhausted, C25 - CD8 T Central Memory, C45 - Naive B, C50 - Transitional B, C57 - Classical Monocyte). In total, patient-specific clusters amounted to 40,181 cells, or 8.9% of the total dataset, confirming that the majority of cell states were shared among disease states and patients.

We annotated AML and CML leukemia cells based on a combination of gene expression profiling (**Figure S1F**)^39^, detection of chromosome Y genes for sex-mismatched (i.e. female donor into male recipient or male donor into female recipient) donor/recipient pairs (**Figure S1G**), and copy number variant (CNV) detection in scRNA-seq data inferred using Numbat (**Figure S1H,I**; **Methods**).^40,41^ Orthogonal validation of AML donor/recipient pairs was performed with Vireo,^42^ using individual DLI product as a reference for donor single nucleotide polymorphisms (SNPs, **Figure S1H**). To enable analysis of shared phenotypes in cases with patient-specific leukemic clusters, we combined myeloid clusters into 7 metaclusters (MC), which were then sub-clustered based on donor or recipient origin and assigned as ‘MC1-7-leukemia’ for recipient-derived cells or ‘MC1-7’ for donor-derived cells (**Figure S1J, Methods**).

Biologic differences between AML and CML cells were preserved in the integrated atlas, revealing distinct clustering of AML and CML leukemia cells, and healthy progenitor cells (**Figures 1C**, **Table S3**). To characterize dominant trajectories of myeloid cell state transition in the integrated atlas, we applied Decipher^43^ (**Methods**), our recently developed tool for studying derailed cell states in disease, which revealed 3 continuous trajectories corresponding to shared myeloid development and disease-specific states. While a normal myeloid differentiation trajectory was observed in both sets of specimens (characterized by high expression of genes consistent with normal myeloid development *VCAN, FCN1, CD14, S100A8, S100A9, S100A12*),^44,45^ we detected divergent CML and AML leukemic evolutionary paths. The AML derailment trajectory was marked by high expression of *CXCL8, RACK1, B2M, CD74, SAT1, TRIM8,* and *IL18,* consistent with previously reported markers of leukemogenesis in AML.^46–51^ In contrast, CML clusters exhibited high expression of genes previously identified as upregulated in this leukemia (*KCNH2, HIST1H4C, HSPA8, RPLP0*, and *C1QBP*).^52,53^ The non-leukemia clusters were overall evenly distributed by response status, disease, and by donor/recipient status (**Figures 1D-E and S1H**), indicating robust correction for batch and library size. Expression of lineage-defining genes (e.g. *CD34* for HSC/leukemia, *CD3E* for T cells, *CD14* for myeloid, *NCAM1* for NK, *CD19* for B cells) confirmed the annotation of major cell types (**Figures 1B and S1K**). While most clusters (n=36, 67%) were composed of cells from both AML and CML patients, 13% (n=7) were dominated by cells originating from AML patients and 19% (n=10) were primarily composed of cells from CML patients after normalizing for differences in total cell number between AML and CML patients (**Figure 1F**). For example, T-cell C40, AML-enriched C42 and C31, and HSC C46 were found primarily in AML patient samples whereas CML-enriched clusters (C7, C33, C47, and C56), B cell C50, and myeloid C51, C54, and C57 were predominantly composed of cells from CML samples. Altogether, construction of this cell state atlas revealed distinct disease phenotypic trajectories and identified specific immune cell populations of interest, some of which were shared while others were unique to each disease.

### Response to DLI in AML entails a cascade of leukemia-immune cell interactions involving ZNF683^Hi^ CD8+ cytotoxic T cells

To systematically assess the associations of the diverse identified cell subsets with response to DLI, we delineated individual cluster composition of marrow-derived cells by disease (AML vs CML), timepoint (pre-DLI vs post-DLI) and response (R vs NR) (**Table S3**). We primarily focused our attention on discerning whether we could detect expansion post-DLI, raising the possibility that these were cellular mediators of response or resistance, although we also evaluated if there was enrichment in either R or NR pre-DLI, that could suggest possible associations with prediction of treatment response. Of 46 clusters with at least 15% of cells detected in CML, we observed only one such expanding cell population: T cell C2, which trended toward expansion in R but not NR (p=0.06, **Figures 2A** and **S2A**), and expressed *CD4, SELL, CCR7, TCF7,* and *IL7R* (**Figure S1L-M**), concordant with our prior detection of an expanding T_PE_ cluster in CML DLI responders.^5^ This cluster, however, was not expanded in AML-R, suggesting involvement of alternate cell types in GvL response in AML.

**Figure 2.**
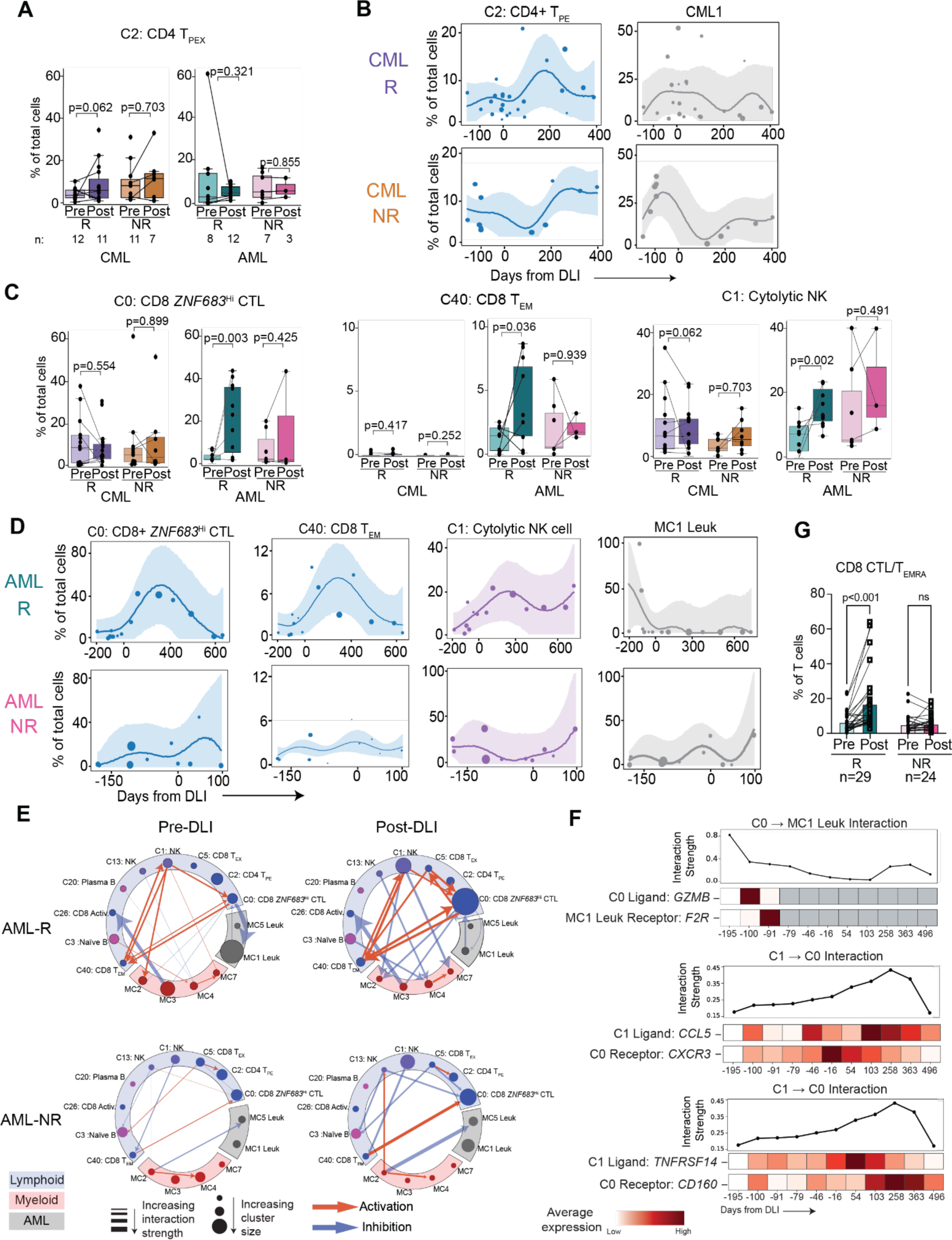
Temporal dynamics and interactions of marrow-infiltrating Immune cell states. **A.** Percentage of C2: CD4 T_PE_ of all cells per patient demonstrating a trend toward expansion in CML-R but not CML-NR (*left*), nor in AML (*right*). **B.** DIISCO prediction for % total marrow cells comprised of the C2 (CD4 T_PE_) and CML1 clusters over time for CML-R (*top*) and CML-NR (*bottom*). **C.** Expansion of C0: CD8 *ZNF683*^Hi^ CTLs (*left*), C40: CD8 T_EM_ (*middle*), and C1: Cytolytic NK cells (denoted as % total marrow cells comprised of a cluster per patient), in AML-R but not AML-NR, nor for CML. Numbers of patient samples evaluated are the identical to panel (A). **D.** DIISCO prediction for C0: CD8 *ZNF683*^Hi^ CTLs, C40: CD8 T_EM_, C1: cytolytic NK cells, and MC1 leukemia clusters for AML-R (*top*) and AML-NR (*bottom*). **E.** DIISCO predicts a coordinated network of interactions between C0: CD8 *ZNF683*^Hi^ CTLs, C40: CD8 T_EM_, and C1: cytolytic NK cells, with additional interactions with C0: CD8 *ZNF683*^Hi^ CTLs and MC1 leukemia cells pre-DLI (*top left*) that strengthen post-DLI, and with further interactions between other lymphoid (blue) and myeloid subsets (red) (*top right*) in AML-R. In contrast, DIISCO predicts few interactions between immune cells or between immune and leukemia (gray) cells in AML-NR pre- or post-DLI (*bottom*). **F.** Interaction between *GZMB* and *F2R* is predicted between CD8 *ZNF683*^Hi^ CTLs and MC1 leukemia cells (*top*). Examples of the top predicted interactions between C1: cytolytic NK cells and C0: CD8 *ZNF683*^Hi^ CTLs include *CCL5:CXCR3* (*middle*) and *TNFRSF14:CD160* (*bottom*). **G.** Percent of CD8 CTL/T_EMRA_ cells of all T cells in peripheral blood of independent post-HSCT relapsed AML patients who received DLI (n=29 responders, 24 nonresponders) by flow cytometry.

In contrast, the most striking expansions in AML-R but not in CML-R occurred in T cell clusters C0 (p=0.003) and C40 (p=0.036), and NK cluster C1 (p=0.002) with a trend toward expansion in T cell C5 (p=0.07), while T cell clusters C21 (p=0.01) and C26 (p=0.001) expanded in AML-NR (**Figures 2C and S2B**). T cell C0 displayed high expression of *CD8A* and the transcription factor *ZNF683*/Hobit, which is increasingly recognized as a mediator of CD8+ T cell cytotoxicity and a marker of tissue resident-memory subsets (**Figure S1L-M**).^54,55^ In addition, this cluster highly expressed various effector genes (*GZMB, GZMH, GNLY, PRF1, B3GAT1*; **Figure S1L**). The T cell cluster C40 was characterized by similar expression of *CD8A* and effector genes (*GZMA, GZMB, GZMH, TBX21, GNLY, B3GAT1*) but with lower expression of *ZNF683* compared to C0 yet still higher than other T cell clusters.

scCITE-seq for C0 and C40 did not demonstrate CD62L expression, and CD45RA expression was higher in C0 compared to C40 (p=0.0004, Kolmogorov Smirnov test; **Figure S1N**) and other T cell clusters, suggesting a phenotypic profile most consistent with a CD8+ T effector memory re-expressing CD45RA (T_EMRA_) profile for C0 and a CD8+ T effector memory (T_EM_) profile for C40. Expanding NK cluster C1 displayed high expression of *B3GAT1, FCGR3A,* and *GZMB*, along with higher expression of CD57 by scCITE-seq compared to other NK clusters, most consistent with a cytolytic phenotype (**Figure S1O,P**).^56^

C0, C1, and C40 did not demonstrate expansion between relapse and post-therapy remission in the chemotherapy only/no DLI controls (**Figure S2C**) and had similar median proportion to that of non-relapse control subjects. To exclude the possibility that the quantified expansion of C0, C40, and C1 post-DLI was predominantly due to higher proportion of leukemia cells in the BME pre-DLI, we computed the proportions of these clusters out of total lymphocytes, excluding the myeloid and leukemic compartment. Reassuringly, we continued to observe a strong signal for expansion particularly for C0 in AML-Rs (p=0.007) compared to NRs (p=0.676) and trends toward expansion of C40 (p=0.17) and C1 (p=0.24) in AML-Rs post-DLI remained (**Figure S2D**). Since the AML-R cohort had a longer duration of followup compared to AML-NR, we asked whether the C0 expansion signal was driven by early or late expansion of this cluster. Analysis restricted to only earliest post-DLI samples for AML-R, obtained within 300 days of DLI re-demonstrated marked expansion of C0 in AML-R compared to NR (p=0.003), supporting the notion that biological differences related to response drive this marked expansion (**Figure S2E**). In investigating enrichments pre-DLI, we only found erythroid (C6: p=0.01, C7: p=0.03) and immature populations (C16: p=0.03, C37: p=0.02) enriched in AML-NR compared to AML-R pre-DLI. Altogether, these analyses support the notion that multiple immune cell subtypes in the AML BME undergo dynamic changes following DLI.

The various dynamic patterns of response might arise from complex interactions and cellular crosstalk within the leukemic BME over time, but current computational methods for inferring cell-cell interactions from scRNA-seq data conventionally generate a static model of predicted interactions. To this end, we applied DIISCO^35^, our recently developed Bayesian model which uses a Gaussian process regression network ^57,58^ to identify possible dynamic networks of cell-cell interactions within the local BME as “*interactomes*” (**Figure S3A**, **Methods**). The modeling of the temporal dynamics of each cell state via a Gaussian process provides sufficient flexibility to integrate single-cell data generated from clinical cohorts which typically entail variable and sparse time-points across patients.^5^ To improve statistical power in resolving temporal patterns and interactions, we constructed an interactome of response for each disease by separately applying DIISCO to the aggregate of 14 AML responder samples in the first 1000 days following DLI and to the aggregate of 24 CML responder samples. Likewise, to achieve disease-specific interactomes of resistance to DLI, we applied DIISCO on all 10 AML non-responder samples and then, to the 11 CML non-responder samples between 200 days pre- and 400 days post-DLI (**Methods**).

In the case of CML, DIISCO identified a central role for expanding CD4+ T_PE_ C2 in responders post-DLI, predicted to inhibit CML leukemia (CML1) and hematopoietic stem cells (HSC) (C15), through coordination with mature NK C13, CD8+ *ZNF683*^Hi^ CTL C0, and plasma B cell C20 (**Figures 2B and S3B-D**). This C2-centralized inhibitory role was not observed in CML nonresponders (**Figure S3D**). Our analysis also unveiled a role for immune cell states beyond lymphocytes in responders, particularly, the prediction of crosstalk between classical monocytes (C4 and C9) with erythroid (C8, C11) and transitional B cells (C23). Predicted ligand:receptor complexes correlated with the inferred interactions between C2 and CML1 included the protein products encoded by *TNF:FAS* and *CD226:NECTIN2*, interactions associated with effective T cell responses. TNF and CD95 (encoded by *FAS*) are part of the pro-apoptotic pathway, potentially supporting a direct role of CD4+ T_PE_ C2 cells in mediating CML cell death.^59,60^ NECTIN2 binds to CD226 on effector T or NK cells to promote T or NK cell-mediated cytotoxicity^61^ (**Figure S3E**). These predicted interaction networks support the central role of CD4+ T_PE_ C2 in mediating GvL response in CML.

In striking contrast to CML, the DIISCO-based analysis of AML-Rs revealed a cascading response after DLI that centered around CD8+ *ZNF683*^HI^ CTL C0, CD8+ T_EM_ C40, cytolytic NK cell C1 and, a CD8 exhausted T cell (T_EX_) population C5 (**Figure 2D and S4A**). As expected, a strong negative interaction between CTL C0 and AML (MC1 leukemia; **Methods**) was observed in responders (**Figure 2E**), consistent with a productive anti-leukemia immune response driving contraction of MC1 leukemia cells. The response interactome model revealed positive interactions among C0, C1 and C40, supporting the notion that these cell subsets formed a coordinated immune network supporting enhanced GvL (**Supplemental Video 1**). To confirm that these predictions were not primarily attributable to an overall increase in immune-infiltration, we re-trained DIISCO on immune clusters only, and found that this coordinated immune network was reproduced (**Figure S4B**), reinforcing the role of C0, C1 and C40 in DLI response in AML.

Since the proportion of C0 CD8+ *ZNF683*^HI^ CTLs varied inversely with proportion of leukemia cells in AML R over time (**Figure 2D**, **Table 1**), we postulated that this cluster might interact with leukemia cells in a manner that would be activating or exhausting in AML-R versus -NR, respectively. In particular, we observed an inhibitory signal from C0 toward AML MC1 through multiple ligand:receptor complexes including proteins encoded by *GZMA:F2R*, an interaction which has previously been shown to promote CD8 T cell cytotoxicity and tumor suppression^62^ (**Figure 2F**). PAR-1, encoded by *F2R*, has been shown to inhibit proliferation of leukemic stem cells in murine models of AML.^63^ Interaction of the proinflammatory chemokine *CCL4* on C0 CD8 CTLs with *SLC7A1* in AML MC1 suggested enhanced T cell infiltration in the AML-R BME.^64^ Indeed, AML patients with higher expression of *CCL4* and other chemoattractant genes have been reported to exhibit improved survival compared to patients with lower expression of these genes, supporting a role for greater immune infiltration and anti-leukemic T cell activity in this context^65^. Additionally, an antigen presenting role for NK cells C1 was suggested by crosstalk with CD8 CTLs C0 (through *HLA-E/F* and *CD8A*). Finally, interaction between cytolytic NK C1 and CD8 CTL C0 was predicted to involve *CD160* and *TNFRSF14* (HVEM), a pathway implicated in controlling the balance of T and NK cell exhaustion and activation and an attractive candidate for novel immune checkpoint strategies.^66^ Intra-marrow cellular interactions in AML-R following DLI were driven predominantly by CD8 CTL C0 (p<0.0001, Mann-Whitney U test) (**Figure S4B**). Beyond lymphoid cells, DIISCO also revealed inhibitory links in Rs between CD8 CTLs C0, NK C1, and myeloid metacluster MC2 which is composed of dendritic cells and classical monocytes (**Figure 2E** **and S4B; Table S3**). Conversely, in AML-NR, DIISCO did not demonstrate significant activating interactions between T cells and other immune cell types post-DLI.

To extend these observations, we asked whether CD8+ *ZNF683*^Hi^ CTLs could be identified as an expanding population in the peripheral blood as well as BM. Indeed, we were reassured by the analysis of subject R3501, for which scRNAseq and scTCR-seq data from matched peripheral blood and bone marrow were available. Comparison of these samples revealed T cells of the same clonotype in both compartments (**Figure S4C**). Moreover, we confirmed the phenotype of the circulating post-DLI peripheral blood T cells as consistent with C0 CD8 CTLs discovered in the BME on the basis of high expression of *ZNF683* and *GZMH*, and a T_EMRA_ profile that included the expression of CD45RA in the absence of CD62L (from scCITE-seq). We therefore evaluated the proportion of circulating CD8+ T_EMRA_ cells in an independent validation cohort of 53 patients similarly treated for post-HSCT relapsed AML with DLI (n=29 R, n=24 NR), on whom multi-parameter flow cytometry of peripheral mononuclear cells had been collected in real-time before and after DLI at our institution (**Table S4**). Strikingly, we detected marked expansion of the CD8+ T_EMRA_/CTL population following DLI infusion in R (fold increase: 2.56 pre to post, p<0.001) but not in NR (fold increase: 1.3, p=0.474) (**Figure 2G**). A similar increase in AML-R post DLI was also observed in a naive B cell population (**Figure S4D**), which we had observed to trend towards expansion in the discovery BM scRNA-seq dataset (C3 in 4 of 5 AML-R; **Figure S2D and S4A**).

This PBMC confirmation of our BM scRNA-seq findings in a larger independent validation cohort particularly supported a key role of expanding CD8 CTLs in effective GvL in AML, as well as other immune cell populations such as B cells.

### Distinct spatial relationships of immune and leukemia cells in the BME of responders compared to nonresponders

We sought to confirm our DIISCO predictions that CD8+ CTL cells function as a central hub for response to DLI in AML using an orthogonal, protein-based detection method. We applied CO-Detection by indEXing (CODEX)^67,68^ with a panel of 38 antibodies (**Key Resources**) to assess the spatial relationships of immune cells at a single-cell resolution on matched samples from 6 pre-DLI (4 R, 3 NR) and 6 post-DLI (2 R, 3 NR) bone marrow core biopsy specimens obtained concomitantly with the aspirates analyzed by scRNA-seq (**Table S1**). Median time from sample acquisition to DLI was similar for R and NR (**Table S1**). Images were processed by aligning between cycles to account for small tissue shifts during staining and image segmentation in patches of size 2048 x 2048 pixels (total of 32 to 169 patches per sample) was performed with DeepCell^69^ (**Methods**). We initially focused on quantifying the proportion of T_EMRA_ cells (expressing CD3, CD57, and Granzyme B) in 7 (5 R, 2 NR) samples which contained at least 50 segmented cells (**Methods**). In this manner, we confirmed T_EMRA_ expansion relative to all cells in R (**Figure 3A; Methods**), thus confirming scRNA-seq findings on a protein marker level.

**Figure 3.**
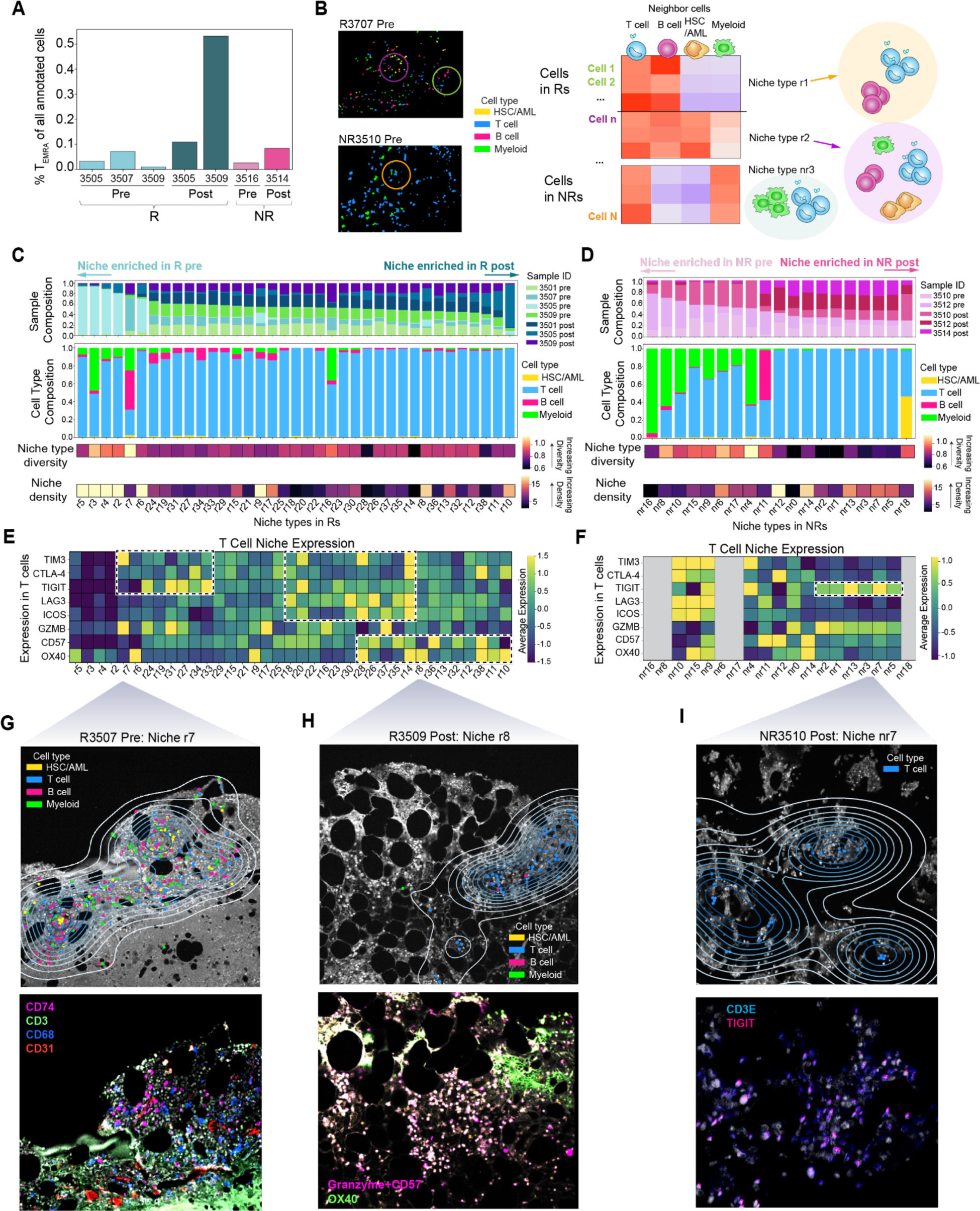
CODEX spatial mapping of coordinated immune cell types in BM core biopsies. **A.** Proportion of T_EMRA_ cells (CD3+ and CD57+ or Granzyme B+) of all cells. **B.** Schema for defining niche types. For each cell, its local spatial neighborhood was defined with a circle of 80 pixels radius, and the number of cells in each major cell type was quantified. Niche types were then defined by clustering all cells across R or NR samples according to their neighborhood composition and density. **C-D.** Niche types identified in Rs (**C**) and NRs (**D**) sorted by ratio of cells pre-DLI to post-DLI. Barplots -distribution of niche types across samples (top) and cell types (middle) after normalizing by total number of cells annotated per sample. Niche type diversity (middle) - entropy of distribution of cell types in each niche. Niche type density (bottom) - cellular density of each niche. **E-F.** Expression of CODEX markers for T cells in each niche identified in Rs (**E**) and NRs (**F**). Grey - niches with <50 cells. Boxes – signatures of exhausted, effector memory, and effector cells. **G.** Example window of a R pre sample (R3507) enriched for niche type r7. Contour line reflects spatial locations with similar distribution of niche type. Smaller distance between contours - steeper change in niche density. Segmented cells are colored by cell type (top) and select markers (bottom). **H.** Same as (**G**) for niche type r8 in an R post sample (R3509). **I.** Same as (**G**) for niche type nr7 in an NR post sample (NR3510). All fluorescence images in **G-I** generated using QuPath^70^ software.

To systematically evaluate the immune networks defining response and resistance, we sought to identify shared spatial neighborhood patterns across samples. Cells were aggregated across all R (4 pre, 3 post) samples, and separately across all NR (2 pre, 3 post) samples, were clustered according to the composition of major cell types in their neighborhood (**Methods**). A neighboring cell was defined as one within 80 pixels (26μm) of the reference cell. Cells without a confident cell type annotation (due to oversaturation of markers) or with >90% of its neighbors unannotated were excluded. Since our marker panel (**Key Resources**) was designed to resolve T cell subsets and other major immune cell types rather than leukemia cells, we focused on defining niches according to immune cell co-localization. Following filtering, 98,130 (75,350 cells for R, 22,780 cells for NR) cells were assigned to one of 4 major cell types (i.e. T or NK cells, B cell, HSC/leukemia, or myeloid cells (**Methods**). The number of cells assigned to each major cell type within its neighborhood was quantified. Recurring colocalization patterns were designated as “*niche types*”, using Phenograph for clustering cells according to neighborhoods. Sample-specific niches composed of >90% cells from a single patient-timepoint were removed (**Figure 3B**; **Methods**). Altogether, we identified 37 distinct niche types in Rs (excluding 2 sample-specific niches) and 19 distinct niche types in NRs (**Table S5**). In certain niches, we observed diverse cell types in their local microenvironment, while others were predominated by a single cell type. Cell density also varied across niches, underscoring the differences in BME based on response (**Figure 3C,D - bottom**).

The R niche types (i.e. r2-r37) were commonly composed of combinations of diverse immune cell populations, including B, T, myeloid, and HSC/leukemia cells across the course of therapy (**Figure 3C**), whereas those identified in NRs (nr1-nr19) lost this diversity post-DLI, as quantified by Shannon entropy (**Figure 3C**,**D**). We additionally observe higher cellular density in the pre-DLI R-enriched niches, compared to NRs, which suggests pre-existing diverse immune cell types in close proximity to be a distinguishing feature of response. This corroborates the notion that coordination of immune networks is closely linked to and may drive GvL response. Additionally, we observed higher abundance of T and B cells dominating niche types enriched in Rs compared to NRs pre-DLI (p=0.004, p=0.04, respectively, Mann-Whitney U test), whereas myeloid cells exhibited higher proportions in NRs vs Rs pre-DLI (p=0.002, Mann-Whitney U test; **Figure 3C,D**). The abundance of T cells further increased in Rs post-DLI compared to pre-DLI in parallel with decreased niche diversity (p=0.0004, Mann-Whitney U test on R pre-enriched vs. R post-enriched T cell proportion; **Figure 3D**), in line with T cell clonal expansion observed with scRNA- and scTCR-seq data.

Deep annotation of T cell phenotypic states across time revealed that the overall expansion of T cells in responders comprised a shift in T cell state from exhausted (expressing TIM3, CTLA-4) pre-DLI to effector (expressing LAG3, Granzyme B, CD57, OX-40) post-DLI (**Figure 3E**). Among post-DLI enriched niches, those of Rs had higher expression of CD57 and OX40 on T cells (Mann-Whitney U test, p<1×10^-15^), while NRs displayed enrichment of niches expressing TIGIT post-DLI (**Figure 3F**). This striking transition to effector memory and cytotoxic T cell states in the R BME complemented the expansion of CD8 CTL C0 and T_EM_ C40 found in our temporal scRNA-seq analysis with DIISCO. Spatial mapping of cell states in niche types enriched in Rs pre-DLI corroborated the co-localization of exhausted T cells with diverse cell types including B cells, myeloid and leukemia cells (**Figures 3G**), while niche types dominating Rs post-DLI include CD8+ CD57+ CTLs co-localized with other cytotoxic and effector memory T cells as well as myeloid cells enriched in dendritic cell marker CD11C (**Figure 3E**,**H**; **Figure S4E**). In contrast, niche types in NRs reflected low diversity and dominance of myeloid cell types (**Figures 3I**).

### C0 defines a CD8+ ZNF683^HI^ T cell population with leukemia specific activity

Our multimodal analyses including scRNA-seq, computational prediction of interactions, spatial phenotyping with CODEX, and validation with flow cytometry of peripheral blood in a larger independent cohort established expansion of C0 CD8 CTLs as a central feature in AML response to DLI. We thus sought to more deeply characterize this population and its differences between AML-R and -NR that might explain mechanisms of response and resistance. Since CD8 CTL C0 expressed transcripts related to T cell effector function (e.g. *GZMA, GZMB, GZMH, IFNG, PRF1*) along with some expression of exhaustion genes (e.g. *TIGIT, GZMK*) (**Figure S1L**), we mapped its position in differentiation and activation trajectories relative to other T cell subtypes. Using Decipher^43^, we identified latent factors denoting transition to exhaustion states (Z8; **Methods; Figure S4F**) and discriminating between CD4 and CD8 subsets (Z9) (**Figure 4A**). T cells within C0 in AML-Rs skewed towards effector states (enriched in *GZMH, CCL5, CXCR4, NKG7, KLRK1*, **Figures 4B-top, S4G-H**) while that of AML-NRs skewed towards terminal differentiation and exhaustion (enriched in *TIGIT, KLRG1, TCF7*) (distribution shift on Z8, p=2.2×10^-11^, Kolmogorov Smirnov test) (**Figure 4A,B and S4I**).

**Figure 4.**
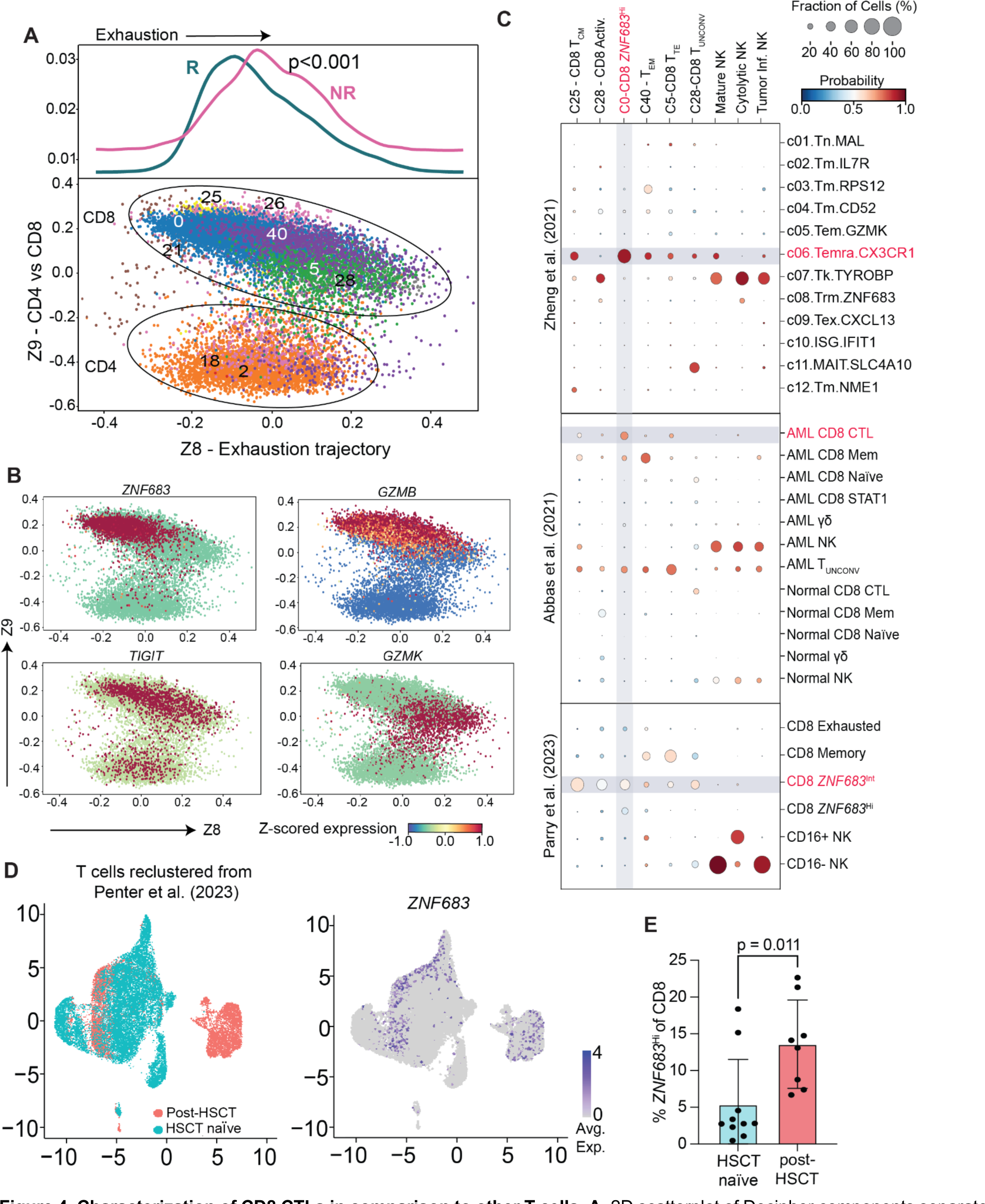
Characterization of CD8 CTLs in comparison to other T cells. **A.** 2D scatterplot of Decipher components separate AML T cell clusters by exhaustion (Z8) and CD4 vs CD8 (Z9) (*bottom*). Distribution of T cells along the exhaustion axis, R vs NR, p=2.2×10^-11^, Kolmogorov Smirnov test. (*top*). **B.** Examples of expression of activation genes (*ZNF683, GZMB*) and exhaustion genes (*TIGIT, GZMK*) along the Z8 axis. **C.** Comparison of gene expression profiles of the CD8 T and NK cell subsets from the current study (columns) versus other published datasets. Dot size - percent of cells in each cluster aligned with the expression profile from the external dataset. Dot color - probability of similar expression to the cluster from the external dataset. **D.** 2D UMAP with cells from HSCT-naive subjects (blue) and post-HSCT subjects (salmon pink) (*left*) from Penter et al (2023). Coloring of UMAP to show expression of *ZNF683* (*right*). **E.** Percent of CD8 *ZNF683*^Hi^ T cells of all CD8 T cells in HSCT-naive (blue) and post-HSCT (salmon) patients.

Indeed, comparison of these T cell states with those found in previous studies of the TME confirmed the CD8 CTL and T_EMRA_ characteristics of C0 (**Figure 4C; Methods**). In one analysis, we compared gene expression signatures of our six CD8+ clusters against the T cell metaclusters from a large scRNA-seq pan-cancer atlas of tumor-infiltrating CD8 T cells curated from 316 donors and 21 cancer types.^71^ C0 most closely correlated with CD8 metacluster signature c06.Temra.CX3CR1 cells (**Figure 4C-top**). A key difference between this metacluster and C0, however, was the high expression of *ZNF683* in the latter.^71^ Second, we evaluated profiles of marrow-derived T and NK cells from normal donors and patients with AML treated with ICB.^72^ C0 most closely aligned with CD8+ CTL populations from AML patient marrow, but not with normal marrow T cells (**Figure 4C-middle**). Finally, we compared our T cell populations with those from a recent study of Richter syndrome, which identified a population of *ZNF683*^Hi^ CD8+ T cells associated with response to PD-1 ICB.^73^ C0 was most transcriptionally similar to the previously reported *ZNF683*-intermediate (*ZNF683*^Int^) CD8+ T cell population, followed by the *ZNF683*^Hi^ CD8+ subset (**Figure 4C-bottom**).

In support of the notion that C0 represents an effector T cell population as opposed to a cytotoxic NK-like population, we compared its expression profile to a population of NK-like CD8+ T cells marked by a distinct gene signature that includes *BCL11B*^Low^ expression, recently described to be expanded by human cytomegalovirus exposure (CMV) with effector function against leukemia cell lines^74^ (**Figure S5A**). While the three NK clusters C1, C13, and C36 aligned closely with the characteristic NK-like genes of this signature, none of our T cell clusters, including C0, expressed the NK-like signature, confirming C0 CD8+ CTLs as distinct from the previously described NK-like CD8 population recognizing CMV.

To define the potential contribution of C0 cells across diverse settings of therapy response in AML, we reanalyzed two published scRNA-seq datasets generated from marrow and blood. First we reclustered marrow-derived T cells collected from post-HSCT-relapsed or HSCT-naive relapsed myelodysplastic syndrome (MDS)/AML patients after treatment with the DNA methyltransferase inhibitor decitabine and ICB ipilimumab and identified four clusters with *ZNF683*^Hi^ expression (cluster ID 2, 3, 4, and 10) similar to C0 (**Figure S5B-D**).^75^ Of note, these cells were more prevalent in post-HSCT patients than in non-HSCT patients (p=0.01, Mann-Whitney test), supporting the notion that they arise in the setting of immune reconstitution following allogeneic stem cell transplantation (**Figure 4D**,**E**). Secondly, we re-analyzed the bone marrow T cells from the ICB-treated AML cohort described above^72^ and identified a subset of *ZNF683*^Hi^ CD8+ T cells, most consistent with C0 (**Figure S5E,F**). We observed a trend toward expansion in R after ICB (p=0.27), but not NR (p=0.85, paired t test; **Figure S5G**).

Since our trajectory analysis of CD8 subsets suggested divergent fates for AML-R vs -NR C0 T cells, we assessed differential gene expression between C0 R and NR cells. Strikingly, *ZNF683* was among the most highly differentially expressed genes (DEGs) in AML-R compared to NR (p<0.0001; **Figure 5A, Table S6**). AML-R C0 cells were marked by higher expression of activation genes (*GZMH, NFKBIA, NKG7, STAT1, JUNB, IFNG).* AML-NRs displayed high expression of *KLRB1* (encoding CD161), recently identified as an inhibitor of tumor-infiltrating lymphocyte (TIL) cytotoxicity in gliomas^76^, as well as multiple genes in the metallothionein pathway (*MT1X, MT2A, MT1F, MT1E*), associated with TIL dysfunction^77^ (**Figure 5A**).

**Figure 5.**
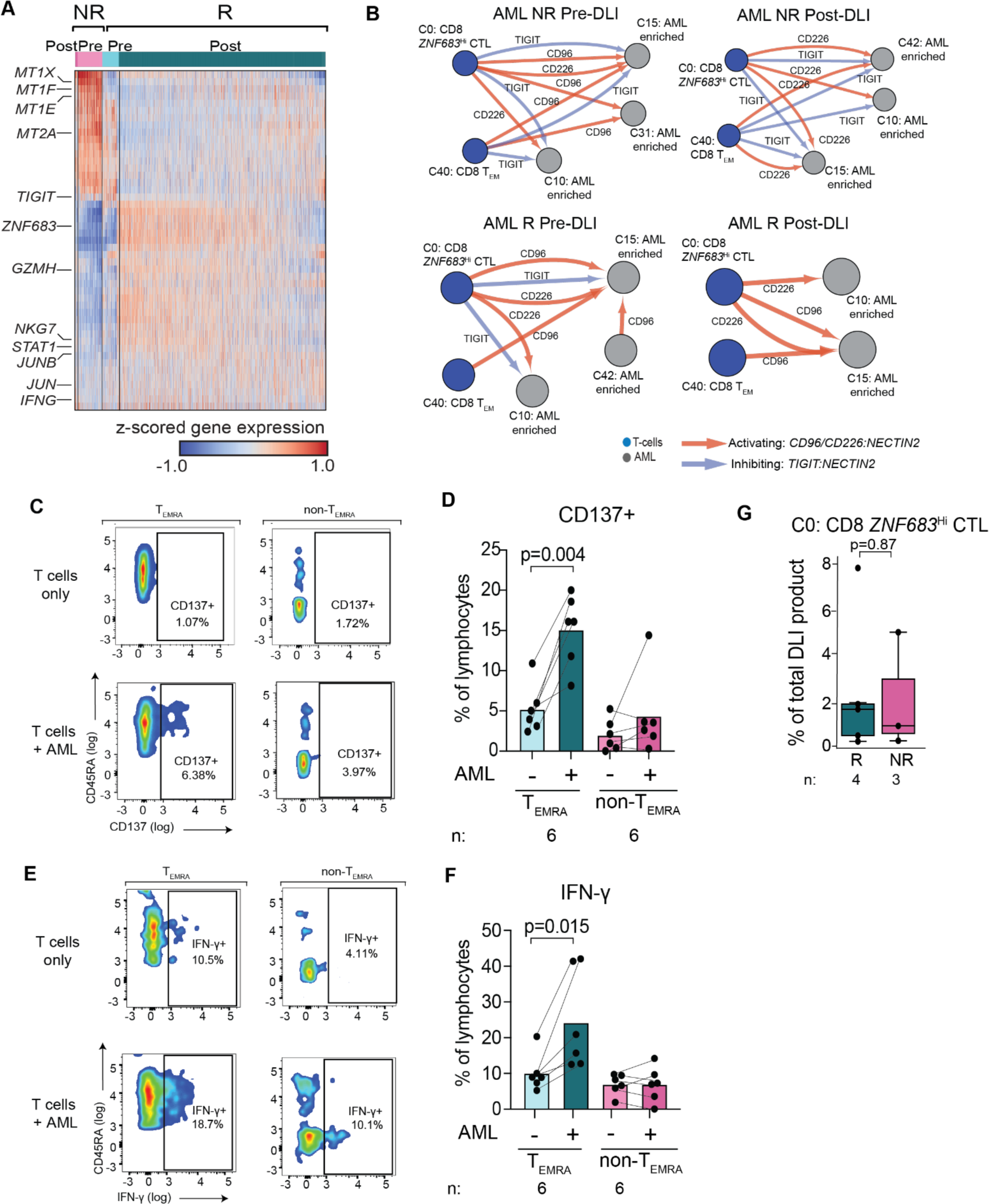
Phenotypic and functional characterization of C0 CD8 *ZNF683*^Hi^ CTLs. **A.** Top differentially expressed genes (DEGs) of AML C0 R (right) versus NR (left) cells. **B.** CellphoneDB networks visualized using CytoScape^84^ highlights interactions between T cell clusters C0 and C40 (blue nodes) with the AML enriched clusters C10, C15, and C42 (grey nodes). **C,E.** Example flow cytometry plots of CD137 (**C**) or IFN-ɣ (**E**) expression in CD8 T_EMRA_ cells alone (*top left*), CD8 non-T_EMRA_ cells alone (*top right*), CD8 T_EMRA_ cells co-cultured with autologous AML cells (*bottom left*), or CD8 non-T_EMRA_ cells co-cultured with autologous AML cells (*bottom right*). **D, F.** Quantification of CD137 expression (**D**) or IFN-ɣ (**F**) across six AML-R patients of CD8 T_EMRA_ or CD8 non-T_EMRA_ cells cultured with or without autologous AML cells. Lines connect samples from the same subject. **G.** Percent C0 of all DLI product cells for R versus NR.

Of the predicted ligand:receptor interactions between C0 and leukemia cells (via CellPhoneDB^78,79^ and CellChat^78^, **Table S7**), that of the exhaustion marker *TIGIT* (encoding TIGIT protein) on CD8+ CTLs and *NECTIN2* (encoding Nectin-2/PVRL2/CD112), a well-described immune checkpoint axis for T and NK cell activation/exhaustion, on AML leukemia cells was amongst the most significant in AML-NR (**Figure 5B**-**top**).^80–82^ TIGIT competes with CD226, an activating co-stimulatory signal, for binding of CD112/CD115 and has stronger affinity, thus controlling the balance of T cell activation or exhaustion. ^61,83^ Accordingly, NECTIN2/PVR interactions with CD226 but not TIGIT were predicted by CellPhoneDB in AML-R, corroborating a more activated profile in R (**Figure 5B**), while AML-NRs were predicted to have TIGIT-NECTIN2/PVR interactions. Thus, CD8+ *ZNF683*^Hi^ CTLs from AML-R are predicted to be more highly activated and cytotoxic after DLI, compared to cells from AML-NR that are more exhausted and dysfunctional, consistent with their potential roles in GvL response and resistance, respectively.

In further support for a likely role of *ZNF683*-expressing CD8+ T cells in mediating anti-leukemia responses, we detected evidence of leukemia-specific reactivity of the CD8+ CTL population. To examine the functional impact of CD8+ CTLs after encounter with leukemia cells from the same patient, we devised an *in vitro* co-culture assay to test leukemia-specific T-cell activation (**Methods**). Strikingly, CD8+ CTLs from 6 AML-Rs expressed higher CD137 (p=0.004) and IFNγ (p=0.015) after co-culture with primary autologous leukemia cells compared to other T cell subsets from the same samples (**Figure 5C**-**F**). CD8+ CTLs cultured in the absence of leukemia cells did not show elevated expression of either CD137 or IFNγ, suggesting antigen-specific functional activation of CD8+ CTLs but not other T cell subsets after encounter with autologous leukemia. These results indicate leukemia-specific activation of CD8+ CTLs, consistent with a potential functional role of these cells in AML DLI responders.

Altogether, these characterizations provide independent support for a likely role of *ZNF683*-expressing CD8+ CTLs in mediating anti-leukemia responses, even across diverse settings of immunotherapy and diseases.

### CD8+ CTLs from the DLI product undergo clonal expansion in AML responders

Given the apparent central role of the C0 CD8+ *ZNF683*^Hi^ CTL population in anti-leukemia response, we hypothesized that the increased proportion of this population in AML-R post-DLI could be due either to higher proportion in the DLI product or greater degree of clonal expansion after infusion, supported by a more immunologically diverse BME as suggested by CODEX. To address the first hypothesis, we analyzed the proportions of cell types within the DLI products of the 4 AML-R and 3 AML-NR from our full dataset that we profiled. No differences in the percentage of C0 cells in the product were observed between R and NR (p=0.88), nor in the other clusters observed to expand in our BM scRNA-seq data (i.e. C1 [p=0.79] and C40 [p=0.19], **Figures 5G** **and S6A**). Moreover, we did not observe differences in proportion of major cell types including total T cells (p=0.43), CD4 T cells (p=0.21), CD8 T cells (p=0.24), total B cells (p=0.66), and NK cells (p=0.82) (**Figure S6B**). In support of the second hypothesis, scTCR-seq analysis of T cell clonotypes within BM and DLI products demonstrated marked T cell clonal expansion primarily in C0, with the majority of these clonally expanded cells deriving from the BM rather than the DLI product (**Figure 6A-C**). Highly expanded clones also expressed more *ZNF683*, while unexpanded clones expressed more *TIGIT*, corroborating our R vs NR DEGs (**Figure 6D**). To determine whether clonal expansion correlated with activation or exhaustion, as suggested by the prior Decipher analysis (**Figure 4A,B**), we analyzed the fold change of genes expressed in C0 CD8 *ZNF683*^Hi^ CTLs versus C5 CD8 T_EX_ and correlation with clonal expansion (**Figure 6E**). Activation genes (e.g. *ZNF683, GZMB, PRF1, FCGR3A, CX3CR1, NKG7*) were more highly expressed in C0, correlating with increased clonal expansion, whereas genes associated with exhaustion (e.g. *TIGIT, LTB, TCF7, GZMK*) were more highly expressed in C5, negatively correlating with clonal expansion.

**Figure 6.**
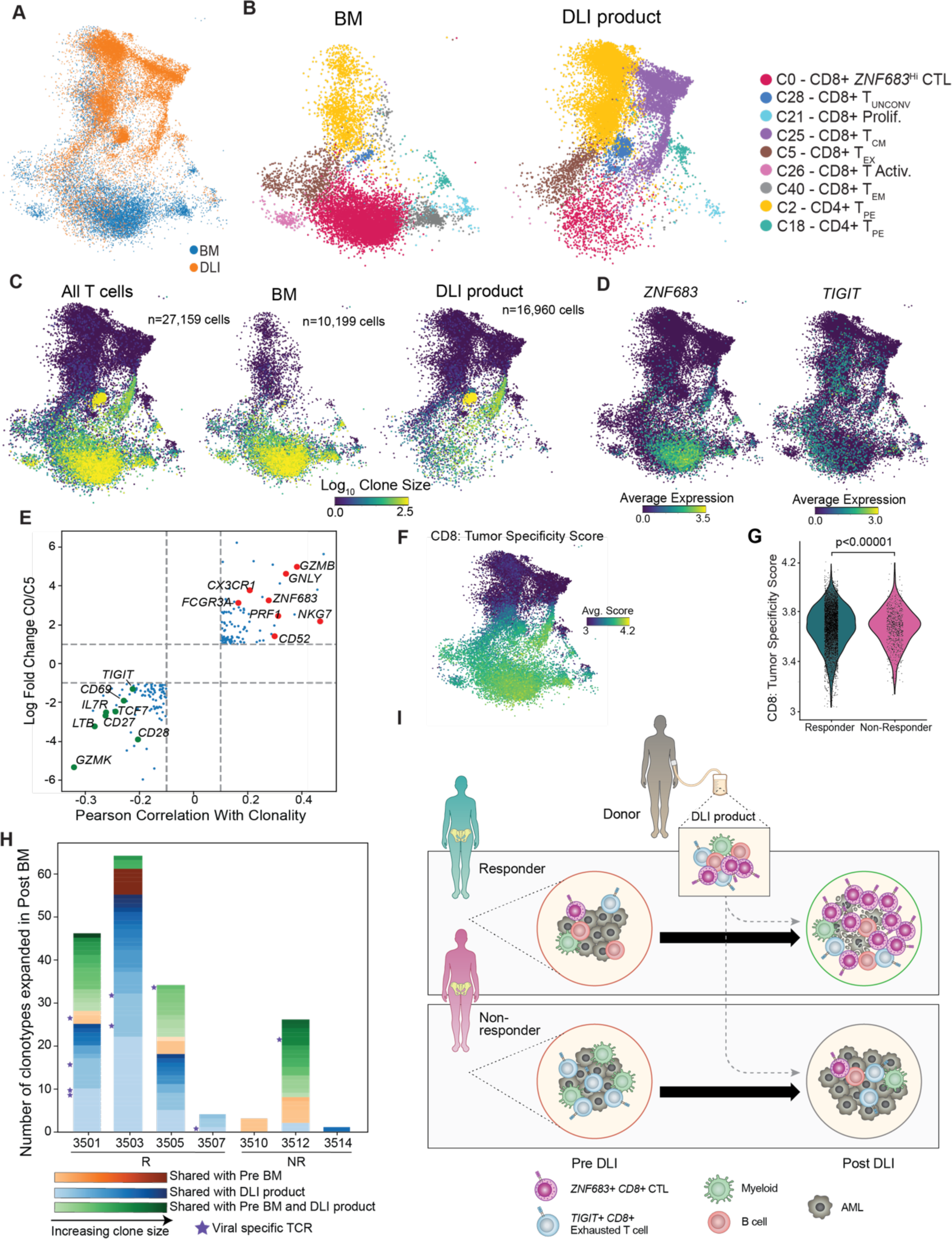
T cell receptor clonal dynamics of C0 CD8 *ZNF683*^Hi^ CTLs. **A.** 2D UMAP of all bone marrow (blue) or DLI product (orange) T cells from AML subjects. **B.** Coloring of 2D UMAPs of BM (left) and DLI product (right) T cells by cluster. **C.** Coloring of 2D UMAP by degree of clonal expansion for each T cell clone for all T cells (n=27,159, *left*), bone marrow T cells (n=10,199, *middle*), and DLI product T cells (n=16,960, *right*). Yellow on color scale indicates greater clonal expansion (measured as Log_10_ clone size), while dark blue indicates clones represented by a single member. **D.** 2D UMAPs demonstrating average expression of *ZNF683* and *TIGIT*. Higher gene expression indicated by green/yellow. **E.** Correlation plot of increasing clonality (x axis) with greater expression of genes in C0 CD8 *ZNF683*^Hi^ CTL versus C5 CD8 T_EX_ cells. **F.** Coloring of T cells by similarity to a previously published CD8 tumor specificity score. Yellow - higher specificity score; blue - lower specificity score. **G.** CD8 tumor specificity score for C0 CD8 *ZNF683*^Hi^ CTLs from R (teal) vs NR (pink). **H.** Bar plot - number of expanded clonotypes shared between DLI product and post-DLI BM (blue) versus those shared between pre-DLI BM and post-DLI BM (red) or found in all three (green) for each subject. Stars indicate clonally expanded TCRs specific to known viral epitopes. **I.** Schema of proposed model.

To assess for viral specificity of clonally expanded T cells rather than leukemia specificity, we compared clonotype sequences to the viral epitope database VDJdb.^85,86^ Only 3.4% of the expanded clones (36 of 1049 clonotypes with size>1) corresponded to known virus-specific TCRs. Of the 36 unique clonotypes identified with known viral-specificity, 17 clonotypes and 119 sequenced TCRs fell within the C0 CD8+ *ZNF683*^hi^ CTL population (**Figure S6C**). Two recent publications established a gene expression signature of CD4+ and CD8+ tumor-specific tumor infiltrating lymphocytes (TILs) in melanoma^87,88^. In comparing these tumor-specific T cell signatures to our T cell populations, clonally expanded C0 CD8+ *ZNF683*^Hi^ CTLs and C21 CD8+ T_PROLIF_ clusters demonstrated greatest similarity to both the CD4+ and CD8+ tumor specificity signatures (**Figures 6F and S6D,E**). C0 cells from AML-R had a higher CD8 tumor specificity score compared to C0 cells from AML-NR (p<0.00001, **Figure 6G**).

We interrogated whether clonal expansion and activation of C0 CD8 *ZNF683*^Hi^ CTLs associated with effective GvL in AML-R was primarily due to expansion of pre-existing clonotypes in the pre-DLI BM versus expansion of cells from the DLI product.^5,21,89^ Some clonotypes were detected to be overlapping between pre-DLI and post-DLI, DLI product and post-DLI, or across all time points/compartments (**Figure S6F,G**). Clonotypes post-DLI were more likely to be shared with the DLI product rather than the pre-DLI BME in 3 of 4 Rs (R3505: p=1.96×10^-5^, R3503: p=4.61×10^-10^, R3501: p=9.5×10^-12^, R3507: p=1; Fisher’s exact test) and 1 of 4 NRs (NR 3512: p=2.3×10^-13^, NR3516: p=1, NR3510: p=1, NR3514: p=1; Fisher’s exact test; **Figures 6H**, **S6H**). Further, higher TCR diversity was observed in the post-DLI BM of Rs compared to NRs (p=0.01, t-test on Gini coefficient; **Figure S6I**). No difference was observed in TCR diversity within DLI products or the pre-DLI BM of AML-R versus AML-NR (p=0.59 and p=0.6, respectively, t test, **Figure S6I**), suggesting that the phenotypic state of T cells and their communication with other immune cells in the BME is the primary driver of C0 CTL activation and clonal expansion post-DLI, rather than differences in the T cell clonality or composition of the DLI product. Thus, our findings support the notion that, while composition of the DLI product is similar between Rs and NRs, DLI infusion in the less exhausted, more immunologically diverse R BME promotes clonal expansion of highly activated effector C0 *ZNF683*^Hi^ CTLs with GvL cytotoxic activity (**Figure 6I**). Conversely, in NRs, infusion into the exhausted, less diverse BME (i.e. *TIGIT*-high) prevents expansion and cytotoxicity of C0 *ZNF683*^Hi^ CTLs, leading to TIL dysfunction therapy resistance.

## Discussion

Highly coordinated immune cell networks with enrichment of activated or cytotoxic T cells have recently been identified as key features associated with response to immunotherapy across a variety of cancer types, largely focused on solid tumor responses to immune checkpoint blockade (ICB).^88,90–93^ In these settings, communities of diverse immune cells, including T and B cells, often accompanied by antigen presenting cells, have been described to aggregate into tertiary lymphoid structures (TLS) or TLS-like neighborhoods,^94^ which are thought to promote T- and B-cell anti-tumor memory and effector activity.^95–102^ Multiple recent studies of the spatial relationships of immune cells in cancer have pointed toward stem-like CD8+ T cells expressing PD-1 and TCF-1/TCF-7 as a hub for anti-tumor immunity, often in complex with T helper and myeloid cell populations, such as dendritic cells.^90,92,103–106^ For myeloid malignancies, however, such immune networks have been only minimally evaluated (for example, in the pediatric AML setting^107^) in part due to the technical obstacles of applying high dimensional sequencing and spatial profiling approaches to marrow tissue that has been conventionally processed using harsh decalcification procedures, though advancing technologies and analytical methods may overcome this challenge.^108–110^ The response of AML to immunotherapies such as ICB have been disappointing,^65,111^ and defining the role of TLS-like structures or other cellular networks could serve to advance our understanding of the critical cellular players needed to coordinate an effective anti-leukemia response. Indeed, in our study, through integrated single cell analysis of marrow biospecimens collected from a deeply clinically annotated cohort of AML immune therapy responders and nonresponders, with incorporation of time-resolved prediction of cell-cell interactions, protein level validations in independent patients, and functional analysis, we gained numerous insights into the relationships between the immunologically diverse BME and response to the archetypal adoptive cellular therapy, DLI.

Our first major insight was the strikingly different organization of cellular populations within the marrow of AML-Rs compared to AML-NRs. The former demonstrated coordinated dynamic expansion among T and NK cells with strong predicted direct interactions with leukemia cells, whereas the latter revealed a picture of relative ’neglect’ from infiltrating immune cell populations, reminiscent of the distinction between ‘hot’ and ‘cold’ tumors that have been previously described in the context of ICB response.^112–116^ Our predictions were confirmed spatially by CODEX, which further revealed co-localization of T cells with B cell populations within immune cell communities in the BME of AML-R, and also a pre-existing preponderance of co-localization with myeloid cells in AML-NRs. These cell population dynamics were further validated through evaluation of a large independent sample cohort by flow cytometry. A growing body of evidence implicate myeloid derived cells with suppressor functions (MDSCs) in limiting effective anti-cancer immune responses across diseases, perhaps mediated through suppression of CD8+ CTL activity.^117–121^ Altogether, our analyses support a central role for a higher diversity of immune cell types that are locally interacting and coordinating with an evolving profile of T cells, along with NK cells for driving GvL in AML.

Even as we observed multiple immune cell populations creating a network of activity, these appeared to converge upon a distinct CD8+ CTL population as the key feature distinguishing AML-Rs. Thus, our second major insight was the discovery of a CD8+ *ZNF683*^Hi^ CTL subset as a central hub associated with GvL response in AML. Notably, we confirmed that co-culture of this cell population with matched primary autologous AML cells *in vitro* led to marked upregulation of IFN-γ and CD137, supporting their role in direct recognition and activation in response to leukemia cell encounter. Our work aligns with other recent studies that together point to a role for *ZNF683*-expressing CD8+ T cells in response to immunotherapy in blood malignancies and solid tumors,^73,122^ as well as in broader antitumor responses.^122–130^ Conversely, we found that AML-NR CD8+ CTLs were characterized by relatively lower expression of *ZNF683* and high expression of the exhaustion and terminal effector markers *TIGIT* and *KLRG1*, suggesting an exhausted, less diverse BME as a possible contributor of resistance to GvL.

Third, our work demonstrated that the C0 population of leukemia-reactive T cells in AML-Rs originated from the DLI product, based on TCR clone tracking. The cellular composition of the DLI product did not differ between AML-Rs and AML-NRs. We therefore inferred that while the major cellular effectors of GvL (the C0 population) in AML arise from the DLI product, their subsequent clonal activation, expansion and sustained immune coordination in Rs depends on support by a recipient BME that is more diverse and less immune-inhibitory than in AML-NRs. Notably, our results raise the tantalizing notion of the existence of disease-specific mechanisms by which GvL is mediated, and hence the possibility of future tailoring of adoptive cellular therapy based on disease context. This is because the scenario we observed in our AML cohort stands in marked contrast to that which we previously observed in DLI for treatment of relapsed CML^5^, where DLI did not directly supply anti-leukemia cytotoxic effectors, but rather provided ‘immunologic help’ to pre-existing marrow-resident exhausted T cells. The caveats to arriving at such a conclusion, however, are the differences in the composition of DLI products between our current analysis cohorts. The CML cohort was treated on a clinical trial involving CD8+ T cell depletion prior to infusion,^33^ whereas the AML cohort was treated with conventional (i.e. unmanipulated, CD8+ replete) DLI. Though 80% of CML patients in this trial achieved response, suggesting DLI product CD8+ T cells are dispensable for GvL in CML, confirmation of the roles of distinct T cell subsets in mediating response for each disease group awaits comparison with more comparable datasets.

Our work presents numerous opportunities for translational impact while also providing a map for intriguing directions of future inquiry. First, our finding of clear differences in the BME of AML-Rs and AML-NRs points to the potential for using this knowledge to devise a prediction schema for identifying outcome to DLI therapy. Certainly, patients predicted to not respond to DLI could benefit from earlier decision-making to select alternative treatment approaches and for eschewing potential risk of immune toxicities that are known to be associated with DLI. Development of such a prediction schema would benefit from identifying the upstream regulators or determinants of such differences in the BME of these patients, as well as further functional interrogation into the nature of these immune networks to unlock mechanisms of action by which non-T cells contribute to GvL. Second, we identified various axes of interaction among cell populations within the BME that could be exploited for therapeutic benefit. For example, we identified high *TIGIT* expression as associated with GvL resistance, perhaps through interactions with the protein products of *PVR* or *NECTIN2* on leukemia cells. The CD226-TIGIT-PVR/NECTIN axis is a well-established immune checkpoint pathway responsible for fine-tuning T cell activation or exhaustion.^80,83,131,132^ Possibly, given the relative resistance of AML to conventional ICB,^65,111,133–135,111^ TIGIT may be a fruitful target for ICB in a subset of AML patients, although definitive experiments are needed to identify the specific patients who may benefit from such a strategy and to further elucidate possible mechanisms of response or resistance. Third, the intrinsic anti-leukemia activity of CD8+ *ZNF683*^Hi^ CTLs make them a highly attractive candidate for cellular therapy for AML. That these cells derive primarily from the DLI product provides the opportunity for graft optimization and engineering through selection and expansion of this population. Given the increasingly evident role for CD8+ *ZNF683*^Hi^ CTLs in association with response to immune therapies in other cancers,^73,122,126^ understanding the mechanisms of antitumor immune activity of this population has broad implications for improving immunotherapy across tumor types. An area of future interest is discerning the antigen specificity of these cells, since the expression of cognate antigen(s) most certainly drove the clonal expansion of this population in AML-Rs. Recent approaches to predict immunogenic epitopes arising in leukemia cells, including not only tumor-associated antigens, but also neoantigens and minor histocompatibility antigens, along with increasingly high throughput means to link TCR specificity with antigen raise the notion that the genomic features of the relapsed leukemia could also contribute to the fate of this T cell population once infused into the recipient.

As the forebear to modern day adoptive cellular therapy, the study of DLI enables the ready ability to address key questions in this rapidly evolving field, namely identifying those cell populations essential to effective anti-tumor activity, defining their interactions with malignant cells and other immune cell populations, and their kinetics over time. As shown in our study, elucidating the mechanism underpinning GvL has broad implications for better understanding the drivers of immunotherapy response in cancer.

## Supporting information

Main and Supplemental Tables

## Acknowledgements

We thank Doreen Hearsey and all staff from the Ted and Eileen Pasquarello Tissue Bank in Hematologic Malignancies for excellent technical support with banking of clinical samples. We thank patients for their generous contribution of research samples for this study. We are thankful to Dana Pe’er, David Knowles, Nicolas Beltran-Velez, Lingting Shi, Michael Pressler, and Mingxuan Zhang for helpful discussions and feedback.

K.M. is supported by the Lubin Family Foundation Scholar Award. S.L. is supported by the National Institutes of Health, National Cancer Institute Research Specialist Award (R50CA251956). C.P. was supported by the Columbia University Kaganov Fellowship. L.P. is a Scholar of the American Society of Hematology, participant in the BIH Charité Digital Clinician Scientist Program funded by the DFG, the Charité – Universitätsmedizin Berlin, and the Berlin Institute of Health at Charité (BIH), is supported by the Max-Eder program of the German Cancer Aid (Deutsche Krebshilfe) and by funding from the Else Kröner-Fresenius-Stiftung (2023_EKEA.102). P.B. is a CPRIT Scholar in Cancer Research and an Andrew Sabin Family Foundation Fellow at The University of Texas MD Anderson Cancer Center and is supported by an Amy Strelzer Manasevit Scholar Award from the Be The Match Foundation and by NCI grant 1K08CA248458-01. C.J.W. is supported in part by the Lavine Family Foundation. C.J.W. and E.A. are supported by Leukemia and Lymphoma Society grant SCOR-22937-22. E.A. is supported by NIH NCI grant R00CA230195 and NHGRI grant R01HG012875.

## Author Contributions

K.M., C.Y.P., E.A., and C.J.W. conceived and supervised the study. K.M., S.L., J.S., W.L., H.L, K.J.L., and P.B. designed and performed experiments. E.A., C.Y.P., and S.M. designed and developed DIISCO. K.M., D.A., C.M., S.K., and S.L.F. designed the CODEX panel and performed experiments. E.A., C.Y.P., Y.J., J.Y.Z., and C.S. performed analysis of CODEX. E.A., C.Y.P, and D.N. developed statistical techniques and interpretation. E.A., C.Y.P., M.B., L.P., J.R.B, and L.R.O. performed single cell and flow cytometry analysis. K.M., C.Y.P., E.A., and C.J.W. analyzed and interpreted results. R.J.S. and J.R. designed and conducted DLI clinical trials and oversaw patient care, contributed samples, and provided clinical data from the BMT repository.

## Methods

### RESOURCE AVAILABILITY

#### Lead Contact

Further information and requests for resources and reagents should be directed to and will be fulfilled by the Lead Contacts Elham Azizi and Catherine J Wu.

#### Materials Availability

All unique/stable reagents generated in this study are available from the Lead Contacts with a completed Materials Transfer Agreement.

#### Data and Code Availability

Single cell transcriptome, CITE and TCR data will be submitted to NCBI’s Database of Genotypes and Phenotype (dbGaP; https://www.ncbi.nlm.nih.gov/gap) and in the National Center for Biotechnology Information’s Gene Expression Omnibus.

#### Code availability

The DIISCO method is available at: https://github.com/azizilab/DIISCO_public. All code used for producing the results and figures of this manuscript is available at https://github.com/azizilab/dli_reproducibility.

## EXPERIMENTAL MODEL AND SUBJECT DETAILS

### Patient/DLI characteristics

The AML patients in this cohort were treated with DLI at Dana-Farber Cancer Institute, Boston, between 2004-2021. All AML patients were treated with conventional CD8-replete DLI as part of standard of care therapy (not on clinical trials). Response was defined as durable morphologic and molecular (i.e. cytogenetic and/or next generation sequencing detection of disease-associated mutations) remission for at least one year after DLI, while nonresponse was defined as lack of post-DLI disease (i.e. % blasts in bone marrow) reduction or disease progression. To ensure the features under study were associated with durable response, patients were not considered for inclusion in the study cohort if they had a short-term response to DLI (i.e. initial reduction in disease burden followed by relapse/progression before 1 year). Two patients who never relapsed were included as non-relapse controls. Pre- and post-chemotherapy treatment samples from three patients who had post-HSCT relapsed AML and entered durable (>1 year) remission with chemotherapy but did not subsequently receive DLI were also included as chemotherapy only/no DLI controls. The presence of acute and chronic GVHD was graded by standard consensus criteria^136^; grades 0 and I acute GvHD were considered clinically equivalent^137^. CML patient characteristics were previously described.^5^

### Sample collection

Bone marrow biopsies and peripheral blood samples were collected pre- and post-DLI from patients consented to a biobanking protocol approved by the Dana-Farber/Harvard Cancer Center IRB. These studies were conducted in accordance with the Declaration of Helsinki. Bone marrow mononuclear cells (BMMCs) or peripheral blood mononuclear cells (PBMCs) were isolated via Ficoll-Hypaque density gradient centrifugation, cryopreserved with 10% dimethyl sulfoxide, and stored in vapor-phase liquid nitrogen until the time of sample processing. For DLI products, 1ml of cryopreserved DLI apheresis product was obtained from QC vials stored at the time of apheresis.

## METHOD DETAILS

### Sample processing and library preparation for scRNA-, scTCR-seq, and scCITE-seq

CML data were obtained from previously described sequencing runs and all cells were analyzed for the current study, whereas we previously reported only the subset of data originating from T cells.^5^ For preparation of AML samples, cryopreserved primary bone marrow mononuclear cells (BMMCs) or peripheral blood mononuclear cells (PBMCs, for the matched sample from R 3501) were thawed on the day of sequencing at 37°C and dispensed into a warmed solution of 10% FBS, 10% DNaseI (StemCell Technologies, cat. No. 07900) in PBS. The cell suspension was centrifuged at 200g for 10 minutes at room temperature. Viable cells were negatively selected using MACS Dead Cell Removal Kit (Miltenyi Biotec, cat. No. 130-090-101). Collected live cells were resuspended in warmed Cell Staining Buffer (BioLegend). Cells were stained with TotalSeq C hashtag antibodies as well as CITE-seq antibodies (**Key Resources**) according to manufacturer instructions. For batch AML1, after washing three times, cells from four samples were combined into one pool (**Table S2, “**Sequencing Run” column) with approximately 20,000 cells from each sample and diluted to a final concentration of 1000 cells/ μl in 0.04% UltraPure BSA (ThermoFisher). Seven total pools were processed in this way. Subsequent samples (i.e. those in batch AML2, DLI products, and controls samples [**Table S2**]) were run individually. Cells were then taken immediately for scRNA-seq, scTCR-seq, and scCITE-seq. Approximately 50,000 cells from each of the pooled samples or 10,000 cells from each of the individual samples were loaded into one lane of a 10x Genomics Chromium^TM^ instrument (10x Genomics) according to the manufacturer’s instructions. The scRNA-seq libraries were processed using Chromium Single Cell 5’ Library & Gel Bead v2 Kit (10x Genomics). Coupled scTCR-seq libraries were obtained using Chromium^TM^ single cell V(D)J enrichment kit (human T cell) (10x Genomics), and coupled scCITE-seq libraries were obtained using Chromium^TM^ single cell Feature Barcode kit. Quality control for amplified cDNA libraries and final sequencing libraries were performed using Bioanalyzer High Sensitivity DNA Kit (Agilent). scRNAseq, scTCRseq, and scCITE-seq libraries were normalized to 4nM concentration and pooled in a volume ratio of 4:1. The pooled libraries were sequenced on an Illumina NovaSeq S4 platform. The sequencing parameters were: Read 1 of 26bp, Read 2 of 90bp, Index 1 of 10bp and Index 2 of 10bp. The sequencing data were demultiplexed and processed as described below.

### PhenoCycler (CODEX) Formalin-Fixed Paraffin Embedded (FFPE) staining

22 mm^2^ square glass coverslips (Electron Microscopy Sciences, 722204-01) were treated by immersing in poly-L-lysine solution (Sigma-Aldrich, P8920) for a minimum of 24 hours and a maximum of 4 days. FFPE tissue samples were sliced onto coverslips and stained following manufacturer protocols (Akoya Biosciences). Briefly, 5 μm sections were cut onto the coated coverslips and baked for 30-60 minutes at 55-60°C. Sections were cooled to RT for 1-2 minutes before deparaffinization and rehydration by successive 5 minute washes in order as follows: 2x in xylene; 2x in 100% ethanol; 1x each in 90%, 70%, 50%, and 30% ethanol; and 2x in ddH_2_O. Sections were moved to 1x dilution of Citrate Buffer (Sigma-Aldrich, C9999) in ddH_2_O, and antigen retrieval was performed in a Tinto Retriever Pressure Cooker (BioSB, BSB 7008) at a high temperature and high pressure protocol (114°C-121°C) for 20 minutes. Sections were subsequently washed in ddH_2_O before being left to incubate in deionized water at room temperature for 10 minutes. Samples were washed twice in Hydration Buffer (Akoya) for two minutes each, then left to incubate in Staining Buffer (Akoya) at room temperature for 20-30 minutes while the antibody cocktail (**Key Resources**) was prepared. A 60 μL/sample of antibody cocktail was prepared according to manufacturer instructions, with primary antibody dilutions of 1:200. A liquid blocker pen was used on the samples to accommodate this reduced volume, and samples were subsequently covered with the antibody cocktail and left to incubate for 12 to 18 hours at 4°C in a humidity chamber. Sections were then washed twice in Staining Buffer for 2 minutes, and subsequently fixed with a mixture of 1.6% PFA in Storage Buffer (Akoya) for 10 minutes. Sections were briefly washed three times in 1x PBS, incubated in ice-cold methanol for 5 minutes, and washed three times in 1x PBS again. Each section was covered with a 190 μL of a mixture of 20 μL Fixative Reagent (Akoya) in 1 mL 1x PBS to incubate at RT for 20 minutes. Sections were subsequently washed three times in 1x PBS and stored in Storage Buffer at 4°C until the assay was ready to be run. Maximum storage time of coverslips did not exceed 2 weeks. A detailed version of the protocol is located at dx.doi.org/10.17504/protocols.io.brznm75e

### Phenocycler/CODEX Assay

A 96-well plate of reporters with DAPI Nuclear Stain (Akoya) was prepared. Samples were loaded in the PhenoCycler according to manufacturer instructions. The section was manually stained with a 1:1000 dilution of Nuclear Stain in 1x CODEX Buffer (Akoya) for 3 minutes before proceeding with the imaging protocol. Imaging was performed on an Andor Dragonfly 200 Spinning Disk Confocal Microscope with a Plan Apochromat 20x/0.75 NA objective (Nikon, MRD00205). To coordinate between the CODEX Instrument Manager and Andor’s Fusion microscope software, the Dragonfly REST-API was used to execute a custom Python triggering script. A pentaband dichroic mirror (405/488/561/640/730, Andor) was used to split emission (“em”) from excitation (“ex”) light with no additional excitation filters. For each round of staining, four channels of confocal imaging were acquired (405 nanometers [nm] ex, 445/46 nm em; 488 nm ex, 538/20 em; 461 nm ex, 594/43 nm em; 640 nm ex, 698/77 nm em) followed by two channels of widefield imaging (405 nm ex, 445/46 nm em; 730 nm ex, 809/90 nm em). Tile scan regions were defined with 10% overlap, and the Nikon Perfect Focus System was used to maintain tissue focus. Images from a single Z plane across the sample were acquired with no binning on an Andor Zyla 4.2 camera. A typical imaging run lasted 48-72 hours.

### Peripheral Blood Expansion Cohort Flow Cytometry Validation

Peripheral blood mononuclear cells (PBMCs) were prospectively collected at pre-specified time points post-HSCT for patients treated on standard of care treatment protocols and consented to the aforementioned biobanking protocol at DFCI. From this patient and sample database, a cohort of 53 patients clinically similar to the scRNA-seq discovery cohort was collected: patients with post-HSCT relapsed AML treated with DLI. Response and nonresponse were defined as in the scRNA-seq discovery cohort. Twenty-nine responders and 24 nonresponders were identified who had flow cytometry data available both at the time of relapse and post-DLI. For responders, pre-DLI samples were analyzed at a median of 55 days prior to DLI (range: day -294 to 0, relative to DLI) and 120 days post-DLI (range: day +14 to +495 relative to DLI). For nonresponders, pre-DLI samples were analyzed at a median of 86 days prior to DLI (range: day -307 to 0, relative to DLI) and 45 days post-DLI (range: day +0 to +317 relative to DLI). Fresh PBMCs were stained with a cocktail of antibodies designed for annotation of T and B cell subsets (**Figure S7A,B**), as described previously,^138^ and data was acquired by multiparameter flow cytometry (LSR Fortessa II) then analyzed using FlowJo (Tree Star). CD8+ TEMRA cells were identified as the CD3+CD8+CD45RO-CD62L-population.

### *In vitro* experiments

From 6 AML DLI patients (2 from the scRNA-seq discovery cohort and 4 from the flow cytometry validation cohort), we obtained peripheral blood T cells at remission (post-DLI). These were cultured with leukemia cells from peripheral blood or bone marrow at the time of relapse after HSCT at an effector to target ratio of 1:1 or T cells alone as a negative control. After 18 hours of *in vitro* coculture, cells were stained for 20 minutes at 4°C in Cell Staining Buffer (BioLegend) with surface antibodies (CD34, CD3, CD4, CD8, CD62L, CD45RA) (**Key Resources**) at a 1:100 dilution. T-cell functional state was assessed by IFNγ catch assay according to manufacturer instructions (Miltenyi). In a separate aliquot of cells from the same well, cells were stained with the same surface antibody cocktail as before along with CD137 to assess for antigen-specific T-cell activation^88,139^. Cells were then washed three times before resuspending in FACS buffer (5% v/v FBS in PBS) and data was acquired using a BD Fortessa II instrument then analyzed using FlowJo (Tree Star) software. Cell subsets were defined as follows: CD8+ T_EMRA_/CTL (CD3+CD8+CD62L-CD45RA+) and CD8+ non-T_EMRA_/CTL (CD3+CD8+CD62L-CD45RA-and CD3+CD8+CD62L+CD45RA+/-) (**Figure S7C**).

## QUANTIFICATION AND STATISTICAL ANALYSIS

### Preprocessing of single-cell RNA, CITE, TCR sequencing data

Quantification of counts was performed using 10X Genomics Cellranger (5.0.0) (**Key Resources**). Quality Control metrics are provided in **Table S2.** There were a total of 12 sequencing runs for AML samples, 7 of which were multiplexed using 2 hashing antibodies per sample, the feature reference sequence of which were then used to demultiplex cells from individual samples/patients in each run. Vireo was applied to demultiplex cells from each of the 4 individuals in the pooled samples, using single nucleotide variants (SNPs) for cell assignment, as well as for determining recipient or donor origin for each cell^42^. DLI product scRNA-seq data was used as “ground truth” for donor-derived sequencing to confirm Vireo assignments, as detailed below. Any cells that were labeled as doublets by the algorithm were removed from analysis. Counts outputs were analyzed using scanpy (v1.9.1)^140^ in AnnData format. All cells with <1000 UMIs were filtered out, leaving 461,940 cells across all datasets for batch correction. CITE-seq protein expression data was normalized using the muon package^141^ for each sequencing pool individually for AML and DLI product. High quality cells were defined based on log_10_ counts of CITE-seq proteins and thresholded between 2.2 and 4 for AML patient samples, and 2.8 and 4 for DLI product samples. Single cell TCR-seq reads were aligned to the GRCh38 reference genome and consensus TCR annotation was performed using Cell Ranger V(D)J (10x Genomics, cell ranger 6.1.2, V(D)J reference `vdj_GRCh38_alts_ensembl-5.0.0`).

### Data integration and batch correction

Due to profiling with different chemistries (3’ vs 5’ for samples with paired scTCR-seq), we performed batch correction on count data using scVI^36,37^ and limiting genes to the top 8000 most highly variable genes (HVGs), with each patient sample added as a separate batch. HVGs were calculated using scanpy’s `highly_variable_genes` function and the ‘`seurat_v3`’ method. The model learned 30 latent components that were used for visualization and clustering. Two patients (5311 and 5322) were sequenced using both 3’ and 5’ chemistries and used to assess batch correction. While cells are grouped together by chemistry before scVI, we observe more overlap between cells of the same patient (despite different chemistry) after integrating with scVI (**Figure S1D**). Immune compartments showed increased mixing on a cluster level, while disease-specific regions were preserved (**Figure 1F**). All samples, including DLI product and controls, were included in batch correction integration. After scVI, data was normalized to median library size and log transformed. In addition to the 10 sample-specific clusters (C10, C42, C46, C54, C56, C18, C25, C45, C50, C57), 6 other clusters were also patient-specific (C21, C28, C30, C43, C48, C51), with >60% of cells coming from a single patient.

### Cell state inference and annotation

Cells were clustered based on inferred scVI latent factors. Clustering was performed with Phenograph (using k=20 KNN) (**Figure 1B**). Forty-three cell states (81%) had good mixing of samples and i.e. had <60% cells from one sample (**Figure S1E**). The threshold for sample-specific clusters was chosen based on the histogram of all sample proportions across all clusters. For proportions of data (AML vs. CML), a threshold of 85% was based on the histogram of distribution of proportions of disease type across all clusters. Due to differences in the number of cells in CML vs. AML, sample proportions were normalized by the number of cells from each disease type (**Figure 1F**). AML-dominant clusters were: 26, 31, 40, 42, 45, 46, 48. CML-dominant clusters were: 7, 16, 33, 37, 47, 50, 51, 54, 56, 57. Four clusters did not have cells from the other disease state but did have cells from control or DLI product samples (47, 50, 57 in CML, 46 in AML).

Cell annotation at the cluster level was performed using differential gene expression analysis and Celltypist^142^. Differentially expressed genes (DEGs) were calculated with the `scanpy.tl.rank_genes_groups` function using the Wilcoxon rank sum test with Benjamini-Hochberg correction. Clusters were first compared to all other clusters to define major cell types and then compared to only clusters from the same cell type for more refined cluster annotation. The same method was used when comparing DEGs within a cluster stratified by response. DEGs were then examined within each major cell type of T, B, and NK cells to manually determine cell subset annotation (**Table S3**).

T cell cluster C0 displayed high expression of *CD8* and differed from other T cell subsets by high expression of *ZNF683*, CD45RA by CITE-seq, and high expression of effector genes (**Figure S1L,M**). We thus designated this cluster as a CD8 *ZNF683*^Hi^ CTL/T_EMRA_ population.^55,143,144^ The remaining T cell populations were characterized as CD4 T_PE_ (clusters 2: *CCR7, TCF7,* and *IL7R*)^5^, CD4 naive (cluster 18: *SELL, CCR7, IL7R*) CD8 exhausted (T_EX_) (cluster 5: *KLRG1, TIGIT, PDCD1, GZMK*),^145,146,147^ CD8 T_CM_ (cluster 25: *CXCR4*),^88,148^ T proliferating (T_PROLIF_) (cluster 21: *MKI67*), and CD8 effector memory (T_EM_) (cluster 40: *CCL5, GZMA, GZMH, TBX21, ID2*),^149^ CD8 activated (cluster 26: *FAS, ENTPD1, IL2RA, TNFRSF4*),^150^ unconventional T cells (i.e. MAIT and ɣδ T cells, cluster 28: *TRGV9*).^151^

B cell clusters 3, 44, and 45 were characterized by expression of *MS4A1, SELL, CR2, PLPP5, IGHD, FCER2*, most consistent with naive B cells^152^, while the remaining clusters included pro-B cell cluster 22 (*CD34, KIT, SOX4, OST4,* and *MME*), pre-B cell cluster 14 (*MME, PAX5,* and *CD24* without *CD34*)^152,153^, transitional B cell clusters 23, 49, and 50 (*MS4A1, FCER2, CD72*), and plasma B cell clusters 20 and 88 *CD19* and *MS4A1* but with high expression of (*JCHAIN, XPB1, IGHG1,* and *IGKC*, without *CD19* and *MS4A1*).^152^ The 3 NK cell clusters included early/mature NK cluster 13 (*SELL, CCR7,* and *NCAM1*),^154^ terminally differentiated/cytolytic NK cluster 1 (low *NCAM1* [and CD56 by CITE-seq), high *FCGR3A* and *GZMB*),^155^ and tumor infiltrating NK cluster 36 (*XCL1, XCL2, CCL3, FOS, AREG*) (**Figure S1O,P**).^156^

Leukemia cell annotation was performed through detection of chromosome Y genes for sex-mismatched donor/recipient pairs, and genetic variant detection in scRNA-seq data inferred using Numbat and Vireo^40,42^ described below.

#### Chromosome Y analysis

CML R5310, CML R5317, AML R3509, AML NR3514 were male patients with female donors. CML R5309, CML NR5318, CML NR5325, CML NR5316, AML R3507 and AML R3503 were female patients with male donors. Patients with a mismatched sex donor accounted for 30.3% of the CML data, accounting for 93,661 total cells. Presence of chromosome Y genes was used in these patients to help validate leukemia cell annotations. Genes used for analysis include: *RPS4Y1, ZFY, LINC00278, PCDH11Y, DDX3Y, EIF1AY, KDM5D, PCDH11Y, TMSB4Y, TTTY14, USP9Y, UTY, ZFY*. We calculated a nearest neighbor graph for each patient using the binarized count matrices. If a cell had >50% nearest neighbors with positive chromosome Y expression, then positive chromosome Y expression was inferred for that cell. 45,356 cells with chromosome Y expression or inferred chromosome Y expression were indentified. Male patients with female donors had chromosome Y expression enriched in clusters 6, 7, 33, 47, and 56. These clusters are defined as “CML-enriched”. For DIISCO, 6, 7, 47, and 56 are grouped as cluster ‘CML1’ and cluster 33 is labeled CML2.

#### BCR-ABL signature for identifying CML-enriched clusters

We created a BCR-ABL signature using the top 20 DEGs upregulated in BCR-ABL+ HSCs relative to BCR-ABL-HSCs.^39^ DEGs calculated using Mann-Whitney U test to rank genes by p-value. Top genes used in signature include the following: *ABL1, RELL2, RNF111, CBR4, OGT, TPRA1, FEZ2, MIR155HG, ZNF665, ZNF852, HOXB2, FAM98B, PYGL, TXN, RXFP1, SRSF3*.

#### Numbat CNV inference

We applied Numbat, a haplotype-aware CNV caller which infers CNVs from scRNA-seq data, to annotate AML cells.^40^ For each AML sample, the following inputs were provided to Numbat: 1) a gene-by-cell count matrix of candidate AML cells (those in non-lymphocyte clusters), 2) an allele dataframe containing SNPs detected in the candidate cells, and 3) a reference gene-by-cell count matrix containing T, B, and NK cells. The allele dataframe was generated by running a preprocessing script provided by Numbat to count alleles and phase SNPs based on a common SNP VCF and phasing reference panel from the 1000 Genomes project. Numbat successfully detected CNVs in three samples (NR3512_1, NR3512_2, and NR3514_1), and the copy number profiles matched expected CNVs from karyotyping (**Figure S1H,I**). To classify AML cells, we filtered cells which passed a 0.9 probability threshold for the expected CNVs (e.g. P(5- or 17-) >= 0.9 for sample NR3512_1) (**Figure S1I**).

#### Vireo analysis

Vireo^42^ was used for orthogonal validation of donor/recipient pairs and for demultiplexing. Cellsnp^157^ was used to obtain SNPs for donor samples and mixed patient-donor samples. SNPs were detected using `genome1K.phase3.SNP_AF5e2.chr1toX.hg38.vcf.gz` as reference. Donor SNP references were defined based on bam files from DLI product sequencing runs, and used to differentiate donor- and patient-cells in the multiplexed AML samples. The number of expected genotypes were set to n=8 in all pools except for Run 3 Lane 1, which was set to n=6 (**Figure S1H**).

#### Myeloid metacluster identification

For myeloid metacluster identification, all bone marrow AML patient cells from the following clusters were included: 4, 6, 7, 8, 9, 10, 11, 12, 15, 16, 17, 19, 22, 24, 27, 29, 30, 31, 32, 33, 34, 37, 39, 41, 42, 46, 47, 51, 52, 54, 55, 56, 57. Clustering was performed on all cells from these clusters using the inferred scVI latent factors in the global atlas of cells via Phenograph and a kNN graph with k=100 neighbors. Cells were re-embedded in new UMAP^158^ coordinates. Metaclusters were annotated based on DEGs and enrichment of patient-derived cells and cells with inferred CNV from Numbat. We also confirmed annotations with Pearson correlation between overlapping genes in above myeloid clusters to bulk data data from 38 distinct populations of human hematopoietic cells (**Figure S1J**).^159^

#### Doublet removal

To detect doublets, in addition to vireo processing, we used Scrublet^160^ on each individual sample. We then calculated the percentage of identified doublets in each cluster and removed clusters that had >20% doublets. The percentage of detected doublets was consistent with expected rates from 10X Genomics. The following cluster IDs were removed: 32, 34, 35, 43, 53.

#### Trajectory analysis

To define and compare disease-specific trajectories, we ran Decipher^43^ on all bone marrow cells from AML and CML patients, limiting to non-erythroid myeloid clusters by excluding the following cluster IDs: 13, 36, 1, 21, 40, 26, 28, 0, 25, 5, 2, 18, 45, 49, 20, 35, 38, 3, 50, 44, 53, 14, 23, 48, 32, 43, 22, 8, 19, 27, 11, 12, 16, 51. Decipher was configured with 10 latent dimensions, 2 visualization dimensions, and beta = 0.1. The model was trained with a learning rate of 0.01 for 16 epochs. We then sorted cells based on Decipher latent components and to show genes of interest that were found to be most correlated with each trajectory (via Pearson correlation). Gene expression was z-scored and then smoothed for each trajectory using a Gaussian filter. Z7 was most consistent with myeloid maturation, while Z2 was most consistent with CML states and Z4 was most consistent with AML states. (**Figure 1C**).

#### T-cell specific Trajectory analysis

For deeper characterization of T cells in AML, we extracted the AML patient cells from all T cell clusters (cluster IDs: 0, 2, 5, 18, 21, 25, 26, 28, 40). Decipher^43^ was configured with 10 latent dimensions, 2 visualization dimensions, and beta = 0.1. The model was trained with a learning rate of 0.01 for 120 epochs. Z8 was most correlated with exhaustion, while Z9 was most consistent with CD4/CD8 subset separation (**Figures 4A**-**B** **and S4F,H**). Density plots were generated using `scipy.stats.gaussian_kde` and parameter `bw_method = 0.05.`

#### Quantification of cell state enrichments

We quantified patterns of enrichment and expansion or contraction of clusters (**Figures 2A,C and S2A-E**) by calculating the number of cells from each sample in a particular cluster as a proportion of total cells in that sample. The calculation and analysis of cell state proportions is based on the assumption of uniform distribution of cell states in the BME, a unique feature of blood cancers. We then grouped samples into time points and response groups relative to DLI. Using `scipy.stats.ttest_ind` we calculated the following comparisons to determine cluster enrichment by response: 1. AML R PRE vs. AML NR PRE; 2. AML R POST vs. AML NR POST; 3. CML R PRE vs. CML NR PRE; 4. CML R POST vs. CML NR POST. To determine cluster expansion and contraction with treatment we used the same t-test function on the following comparisons: 1. AML R PRE vs. AML R POST; 2. AML NR PRE vs. AML NR POST; 3. CML R PRE vs. CML R POST; 4. CML NR PRE vs. CML NR POST. All comparisons were calculated using a 2-sided t-test.

#### CellPhoneDB

CellPhoneDB^79^ version 2.0.0 was applied to generate interactions in each responder group in each disease separately (AML R PRE, AML R POST, AML NR PRE, AML NR POST, CML R PRE, CML R POST, CML NR PRE, CML NR POST). We limited the analysis to clusters that had more than 80 cells. To filter predicted interactions, we only reported receptor-ligand pairs that had non-zero counts in the respective sender and receiver clusters and p<0.05

#### Identification of dynamic intercellular interactions using DIISCO DIISCO framework overview

DIISCO^35^ is an open-source computational framework for characterizing the temporal dynamics of cell-cell interactions using scRNA-seq data from multiple time points, e.g. from samples pre and post-therapy. Using a probabilistic model, DIISCO combines observed cell type frequencies with prior knowledge of receptor-ligand complexes to infer how cellular interactions are evolving with time. As input, DIISCO requires cell type proportions in each sample: given 𝑁 samples at time points 𝑡_1_, …, 𝑡_*N*_ with 𝐾 predefined cell types based on clustering, we define 𝑦(𝑡_*i*_) as a 𝐾-dimensional vector where the 𝑘-th entry contains the proportion of cells assigned to the 𝑘-th cell type in the sample at time 𝑡_*i*_. DIISCO models the cell type dynamics 𝑦(𝑡_1_), …, 𝑦(𝑡_*N*_) as

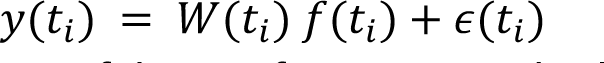

where 𝑓(𝑡_*i*_) is a 𝐾-dimensional vector of latent features such that 𝑓_*k*_(𝑡_*i*_) provides a smoothened approximation to 𝑦_*k*_(𝑡_*i*_). 𝑊(𝑡_*i*_) is a 𝐾 × 𝐾-dimensional matrix which represents temporal interactions and 𝜖(𝑡_*i*_) ∼ 𝑁(0, 𝜎^&^ 𝐼) is Gaussian noise to represent measurement error. DIISCO uses variational inference to infer the parameters 𝑓(𝑡_*i*_) and 𝑊(𝑡_*i*_) which best fit the observed cell type proportions.

The key output of DIISCO is the learned interaction tensor 𝑊. Specifically, 𝑊_*k,k*_′𝑡_*i*_) represents the direction and magnitude of intercellular interaction from cell type 𝑘′ to cell type 𝑘 at time 𝑡_*i*_. 𝑊_*k,k*_′(𝑡_*i*_) > 0 represents an activating interaction, whereas 𝑊_*k,k*_′(𝑡_*i*_) < 0 represents an inhibitory interaction. To avoid learning self-interactions, DIISCO constrains the diagonal terms by setting 𝑊_*i,i*_(𝑡) = 0 in the inference procedure, so that 𝑊_*k,k*_′(𝑡_*i*_) captures the effect of all *other* cell types 𝑘′ ≠ 𝑘 on the dynamics of cell type 𝑘.

To further constrain the solution space of interactions, DIISCO accepts an optional 𝐾 × 𝐾-dimensional binary prior matrix 𝛬, where 𝛬_*k,k*_′ = 1 if cell types 𝑘 and 𝑘′ might interact, and 𝛬_*k,k*_′ = 0 otherwise. In practice, 𝛬 is constructed by evaluating the co-expression of complementary ligand and receptor genes in all 𝐾 × 𝐾 pairwise combinations of sender and receiver cell types (see **DIISCO interaction prior construction).** 𝛬 is used to set the prior distribution of 𝑊(𝑡_*i*_) in DIISCO’s probabilistic model, guiding the inferred interactions to be biologically relevant.

We previously showed the performance of DIISCO on simulated data as well as a controlled experimental setting involving leukemia and CAR-T cell interaction^35^. In this work, we establish the applicability of DIISCO for inferring interactions between a large set of cell states in a complex system such as the BME. In addition, we provide procedures for its application to heterogeneous clinical data with sparse and variable timepoints across patients. We also show the applicability of DIISCO for comparison of malignancies and patient groups, e.g. according to therapy outcome.

### DIISCO data preparation

We first prepared three datasets quantifying cell type proportions over time in the CML and AML samples. The first dataset, CML, contained all clusters in the CML samples. The second dataset, AML(I), contained only healthy immune cell clusters in the AML samples, excluding any Numbat-classified AML cells or recipient myeloid cells. The purpose of this set was to predict interactions according to proportions of cell states of all immune states, thus removing the effect of leukemic regression after therapy. The third dataset, AML(I + L), contained both immune and leukemia clusters in the AML samples, with the myeloid clusters separated into donor and recipient-specific subclusters. Each of these three datasets were further subdivided into cells originating from responder (R) and non-responder (NR) samples. For each of these six datasets, we constructed an interaction prior matrix 𝛬, calculated cell type proportions 𝑦(𝑡_1_), …, 𝑦(𝑡_*N*_), and trained a separate DIISCO model.

### DIISCO interaction prior construction

To leverage prior knowledge on cell signaling in DIISCO, we used a collection of 8,234 literature-curated receptor-ligand (R-L) gene pairs from OmniPath,^161^ a database of molecular biology prior knowledge. Here, we describe the process of using these R-L pairs to construct interaction prior matrices which identify pairs of cell types expected to interact a priori. Note that we created separate interaction prior matrices for each of the six datasets mentioned above to serve as prior knowledge for the respective DIISCO models. First, for each dataset, we filtered out genes with low overall expression, defined as genes with a max log-normalized expression (log(0.1+counts)) across all cells of less than 2 (for the CML dataset), or 1.5 (for the AML(I) and AML(I+L) datasets). Additionally, we filtered out cells in clusters not relevant to the disease type (e.g. filtered out CML clusters for the AML datasets). Of the remaining clusters, we also excluded clusters containing fewer than 1% of the cells in the dataset in order to focus DIISCO on well-represented clusters. For each sample, we identified DEGs in each cluster by performing a Wilcoxon rank-sum test in Scanpy^162^; a gene was deemed a DEG for a cluster if the test had a score > 0 and p-val <= 0.01. Using these sample-level DEGs, we constructed a 𝐾 × 𝐾-dimensional sample-level interaction score matrix 𝑆, where 𝑆_*k,k*_′ is the number of R-L pairs from OmniPath where the ligand is differentially expressed in the source cluster 𝑘′ and the receptor is differentially expressed in the target cluster 𝑘 in that sample. We averaged interaction scores across samples in each dataset to obtain an overall 𝐾 × 𝐾 interaction score matrix 𝑆^<^. We set the diagonal of the interaction score matrix to zero to prevent DIISCO from learning self-interactions between cell types. Finally, we binarized the interaction score matrix by thresholding above the first mode of the interaction scores distribution (e.g. 𝑆^<^ ≥ 5 for the AML(I), R dataset) to create a binary interaction prior matrix 𝛬 for each dataset. 𝛬_*k,k*_′ = 1 indicates that source cell type 𝑘′ might interact with target cell type 𝑘, and 𝛬_*k,k*_′ = 0 indicates otherwise. In the following modeling sections, 𝛬_*k,k*_′ informs the prior distribution of DIISCO’s interaction variables 𝑊_*k,k*_′(𝑡_*i*_), enabling us to encode prior knowledge on cell signaling to guide DIISCO towards learning biologically-relevant interactions.

### DIISCO models for CML datasets

We first filtered out clusters containing fewer than 1% of cells in Rs and NRs. This left behind clusters 0, 1, 11, 12, 13, 14, 15, 16, 17, 2, 20, 23, 24, 3, 4, 5, 8, 9, CML1 in Rs and clusters 0, 1, 10, 11, 12, 15, 16, 2, 22, 25, 3, 30, 37, 4, 41, 5, 8, 9, CML1, CML2 in NRs. Additionally, we filtered out samples before 200 days pre-DLI and after 400 days post-DLI to prioritize interactions more likely associated with treatment response or resistance. This resulted in 24 R samples between -152 days to 392 days from DLI and 11 NR samples between -34 to 295 days from DLI. Cluster proportions 𝑦(𝑡_1_), …, 𝑦(𝑡_*N*_) were calculated for the samples and passed along with the interaction prior matrix 𝛬 as inputs to the DIISCO model, trained separately for R and NR. Model hyperparameters (𝜎_*f*_, 𝑣_*f*_, 𝜏_*f*_, 𝜎_*W*_, 𝑣_*W*_, 𝜏_*W*_, 𝜎_*y*_) were tuned based on visual assessment of predicted vs actual cell type proportions and were kept fixed between the R and NR models, with the exception of the lengthscale hyperparameters 𝜏_*f*_ for 𝑓(𝑡_*i*_) and 𝜏_*W*_ for 𝑊(𝑡_*i*_); these were informed by the distribution of sample timings and set to the mean of all pairwise differences between sample timepoints (𝜏_*f*_ = 𝜏_*W*_ = 150 for Rs and 𝜏_*f*_ = 𝜏_*W*_ = 100 for NRs). The models were trained until convergence of the evidence lower bound (ELBO) objective. By sampling from the trained generative model, we obtained the posterior distributions, for cell type proportions 𝑦(𝑡_*i*_) and temporal interactions 𝑊(𝑡_*i*_) densely sampled over the entire time range of the data. We visualized these predictions using network diagrams and videos (see **DIISCO interaction network visualizations**).

### DIISCO models for AML(I) datasets

We applied a nearly identical modeling pipeline to the AML(I) datasets as we did to the CML datasets above. Filtering out clusters with fewer than 1% of cells resulted in clusters 0, 1, 13, 2, 20, 21, 26, 3, 40, 5, MC1, MC2, MC3, MC4, MC7 in Rs and clusters 0, 1, 13, 14, 2, 23, 26, 3, 38, 40, 5, MC1, MC2, MC3, MC4, MC5, MC6, MC7 in NRs. Filtering out samples after 1000 days post-DLI resulted in 14 R samples between -195 days to 727 days from DLI and 10 NR samples between -182 to 98 days from DLI. Timeframes were selected based on distribution of samples over time-intervals. The lengthscale hyperparameters used were 𝜏_*f*_ = 𝜏_*W*_ = 300 for Rs and 𝜏_*f*_ = 𝜏_*W*_ = 100 for NRs.

### DIISCO models for AML(I + L) datasets

We applied a similar pipeline to the AML(I + L) datasets as we did to the datasets above. Filtering out clusters with fewer than 1% of cells resulted in clusters 0, 1, 13, 2, 20, 26, 3, 40, 5, MC1, MC1_leuk, MC2, MC3, MC4, MC7 in Rs and clusters 0, 1, 13, 14, 2, 23, 3, 40, 5, MC1, MC1_leuk, MC2, MC2_leuk, MC3, MC3_leuk, MC4, MC4_leuk, MC5, MC5_leuk, MC6, MC6_leuk in NRs. Filtering out samples after 1000 days post-DLI resulted in 14 R samples between -195 days to 727 days from DLI and 10 NR samples between -182 to 98 days from DLI. The lengthscale hyperparameters used were 𝜏_*f*_ = 𝜏_*W*_ = 300 for Rs and 𝜏_*f*_ = 𝜏_*W*_ = 100 for NRs.

#### C0 interactome hub

To test how integral C0 was in the entire DIISCO network, we compared the absolute value of W (as interaction strengths) for all links to or from C0 to links not involving C0 in AML responders post-DLI. We tested significance using a Mann Whitney U-test.

### DIISCO interaction network visualizations

To visualize DIISCO’s predicted cell type proportions 𝑦(𝑡_*i*_) and temporal interactions 𝑊(𝑡_*i*_), we created interaction network diagrams using Cytoscape’s py4cytoscape Python package.^84^ Before using Cytoscape, we applied several filters to highlight salient and high-confidence interactions in the visualizations. First, we created a *sustained* filter which selected cell types 𝑘, 𝑘′ for which |𝑊_*k,k*_′(𝑡_*i*_)| was *on average* greater than a threshold during the post-DLI time period (i.e. a sustained, strong interaction post-DLI). Second, we created a *transient* filter which selected cell types 𝑘, 𝑘′ for which |𝑊_*k,k*_′(𝑡_*i*_)| was at *some point* greater than a threshold during the post-DLI time period (i.e. a transient, strong interaction post-DLI). Third, we created a *confident* filter which selected cell types 𝑘, 𝑘′ such that the mean over standard deviation of the predicted cell type proportions for both cell types were greater than a threshold (i.e. cell types for which DIISCO made high-confidence predictions). The filters were combined to select cell type pairs 𝑘, 𝑘′ which satisfied the following rule: ((*sustained* OR *transient*) AND *confident*); only interactions between pairs which passed this rule were shown in the network diagrams. The thresholds used for the filters were tuned to display up to 20 interactions in each R network diagram; the same filter thresholds were applied to the NR diagram to allow for faithful comparison of the R and NR networks for a disease type. For each of the six datasets, we used py4cytoscape to generate an average pre-DLI network, an average post-DLI network, and a video showing the network’s evolution over time. The network’s nodes represent cell types and edges represent interactions between cell types. Node size is proportional to the cell type’s predicted proportion 𝑦_*k*_(𝑡_*i*_). Edge width is proportional to the magnitude of the predicted interaction |𝑊_*k,k*_′(𝑡_*i*_)|, and edge color is based on the sign of the predicted interaction (activating/inhibitory). In the average pre and post-DLI network diagrams, the node sizes, edge widths, and edge colors reflect the average 𝑦(𝑡_*i*_) and 𝑊(𝑡_*i*_) for the corresponding time periods. To generate the interaction network videos, we created network diagrams for evenly-spaced time points over the entire time range and stitched the images into a video using the ffmpeg video processing utility (**Supplemental Video 1**).

### DIISCO receptor-ligand predictions

We identified R-L gene pairs potentially mediating the learned interactions in 𝑊(𝑡_*i*_) by correlating the interactions against R-L gene expression over time. First, we filtered out lowly-expressed genes and underrepresented clusters using the same filters as described in the interaction prior construction step above. For each cell type pair (𝑘, 𝑘′) and each receptor-ligand gene pair (𝑅, 𝐿) from OmniPath where 𝑅 and 𝐿 are differentially-expressed in 𝑘 and 𝑘′ respectively, we calculated correlations between |𝑊_*k,k*_′(𝑡_*i*_)| and 𝑅 expression in cell type 𝑘 over time, as well as between |𝑊_*k,k*_′(𝑡_*i*_)| and 𝐿 expression in cell type 𝑘′ over time. To smoothen the sparse and noisy gene expressions over time, we applied a moving average with a window-length of 2 time-points to the 𝑅 and 𝐿 expressions before computing correlations. Additionally, since 𝑅 and 𝐿 expressions were only observed at discrete sample timepoints 𝑡_1_, …, 𝑡_*N*_, whereas 𝑊_*k,k*_′(𝑡_*i*_) was sampled from the trained DIISCO model at higher-frequency intervals 𝑡_*i*_ ∈ [𝑡_1_, 𝑡_*N*_], we matched each 𝑡_*i*_ to the closest-observed sample in 𝑡_1_, …, 𝑡_*N*_to line up the interactions with 𝑅 and 𝐿 expressions before computing the temporal correlations. After computing correlations, we ranked the R-L pairs by the average of their receptor-interaction and ligand-interaction correlations, and we visualized the top R-L pairs using heatmaps and time series visualizations. This process was repeated for each cell type pair in each dataset.

### Analysis of CODEX spatial data

#### CODEX overview

We imaged 7 responder samples (4 pre-DLI, 3 post-DLI) from 4 R patients, and 5 non-responder samples (2 pre-DLI and 3 post-DLI) from 4 NR patients (Table S1). 4 total samples (2 R, 2 NR) were imaged with only confocal microscope while all other samples were imaged with widefield as well, to better capture low signal in channel 3 (750nm). Each sample was imaged over 18 cycles, with 33 markers across 3 channels, excluding DAPI which was used in channel 0 (405nm) for every cycle. Channel 1 (561nm) markers include: CD14, CD34, Granzyme-B, CD44, CD38, CD21, MPO, CD19, TIGIT, CD8, Ki67. Channel 2 (647nm) markers include: OX40 (TNFRSF4), CD11A, CD152, S100A4, CD74, CD11C, CD68, ICOS, CD4, FOXP3, CD3E, LAG3, PDL1, CD34, TIM3, CD11B. Channel 3 (750nm) markers include: Galectin-9, CD57, CD31, CD47, CD20, Vimentin.

#### CODEX image alignment

Images from samples R3505_post, R3509_post, and R3507_pre were aligned using antspyx^163^ by aligning all DAPI images from cycles 2-18 with DAPI cycle1. For each cycle, alignment generated a transformation that was then applied to all other channels. Each image was saved as a tiff file for downstream processing. Illumination correction for sample R3505_pre was implemented through BaSiCPy.^164^ The corrector was initialized using default basicpy.BaSiC() and applied to the input tile. The corrected image is then clipped to be within a lower threshold and an upper threshold of 5th and 99th percentile of the image’s intensity values, respectively. Finally, the clipped image is normalized from 0 to 1 and the tiles are patched together.

#### Image Registration

Image registration was performed on paired Cycle Fluorescence (CF) and Widefield (WF) imaging platforms from R3509_pre due to offset coordinates between the two sets. For each cycle, we registered WF images (“moving”) against their CF pairs (“fixed”). Panoramic images I^CF^ ∈ ℝ^C×Y×X^, I^WF^ ∈ ℝ^C’×Y’×X’^ were reconstructed from non-overlapping Field-of-View (FOV) tiles. We first applied affine transformation on DAPI channels that are shared across platforms. Specifically, a 3×3 transformation matrix M was computed between I^WF^ and I^CF^ via SIFT[1],^165^ which was then applied across WF channels c_1_,…,c_n_ to obtain the warped image I^WFwarped^ ∈ ℝ^C×Y×X^ towards the CF reference. The pipeline was implemented with `opencv-python`.

#### Post-alignment Processing

All samples were normalized using a min-max normalization and clipping pixel values between 0.05 and 0.99 in order to remove super high-intensity pixel outliers. All normalized images were segmented to define cell boundaries using DeepCell^69^ segmentation algorithm. For set up, we combined pixel intensities from all markers and paired that with the DAPI image from cycle 1 in order to run DeepCell Mesmer pretrained model. We removed any windows with fewer than 50 segmented cells. After segmenting all remaining individual windows, we filtered all pixels within the segmented masks below the 85% percentile to balance needing multiple markers to annotate cells key to study while limiting the number of cells expressing markers from multiple distinct cell types. For all remaining non-zero pixels, we re-normalized and squared the values. We then calculated the average pixel intensity for all markers for each segmented cell. This generates a cell X marker matrix for each sample. We then performed quantile normalization across all cells after normalizing each cell by median intensity across all markers. Any markers expressed in a specific cell at a below average threshold (defined for each marker across all cells in that sample) were set to 0 for that cell. Any cells that had positive expression of more than 12 of 34 markers were removed due to oversaturation. Due to sample-specific differences in pixel intensity distribution, all above steps were calculated within each sample separately.

#### CODEX cell type annotation

T/NK cells were annotated based on non-zero expression of at least one of the following markers: CD3E, CD4, CD8, CD57, TIM3, TIGIT, LAG3, CD152, ICOS, ’FOXP3’, ’Granzyme’,’ TNFRSF4/OX40’. Any cells that were also expressing above average expression of B-cell (defined as mean expression of each marker across all spots), myeloid or HSC markers (’CD19’,’CD20’ for B cells, CD14 for myeloid, [’CD34’,’MPO’] for HSC) were defined as mixed and removed. B-cells were then annotated based on non-zero expression of at least one of the following: [’CD19’,’CD20’,’CD21’]. Any cells that were also expressing above average expression of T/NK (defined as mean expression of each marker across all spots) or myeloid markers ([’CD3E’,’CD8’,’CD57’,’TIM3’,’TIGIT’, ’LAG3’, ’CD152’, ’ICOS’, ’FOXP3’,’ Granzyme’] for T/NK cells, CD14 for myeloid, [’CD34’,’MPO’] for HSC) were defined as mixed and removed. Myeloid and HSC cells were annotated similarly. T_EMRA_ cells were annotated based on above average expression of CD3 and either CD57 or Granzyme B, and lack of expression from contradictory markers for different cell types. Only samples with widefield imaging were included in T_EMRA_ annotations. Percentages shown in **Figure 3A** are based on the number of T_EMRA_ detected divided by the total number of annotated cells detected. Due to lack of widefield imaging, the following 6 markers were excluded from analysis for samples R3501_pre, R3501_post, NR3510_pre, and NR5310_post: [’CD20’, ’CD31’, ’CD47’, ’CD57’, ’Galectin-9’, ’Vimentin’]. Due to sample-wide channel failure, the following 16 markers were excluded from analysis for sample NR5312_post: ’CD11A’, ’CD11B’, ’CD11C’, ’CD152’, ’CD3E’, ’CD4’, ’CD45’, ’CD68’, ’CD74’, ’FOXP3’, ’ICOS’, ’LAG3’, ’PDL1’, ’S100A4’, ’TIM3’, ’TNFRSF4/OX40’.

#### Calculating cell neighborhood

For each sample, neighborhoods were calculated by first defining the center of each cell and calculating its distance to all other cells in that sample. A radius of interaction is defined at 80 pixels, and the cell type of any cells within that radius are tallied. Any cell that is not annotated is excluded. Additionally, to avoid bias of the analysis with uncharacterized neighborhoods, we removed any cells whose neighborhood consists of ≥90% of unannotated cells. This creates a cell x celltype matrix, where the major cell types included are T/NK, B, myeloid, and HSC. All samples were then aggregated by response and clustered with KNN using n=300 neighbors and then phenograph to define clusters of neighborhoods as niches. We tested different neighborhood graphs of n=80, 100, 300, and 600 and found similar clustering between 300 and 600. For cells with at least 1 neighbor, 40% of cells in responders were filtered out and 83% of cells in nonresponders were filtered out.

#### Niche Analysis

Response-specific niches ordered based on percent of cells originating from pre-DLI samples. Due to differences in number of cells per sample, proportions were normalized based on sample-specific counts. Cell type composition was calculated based on the number of cells from each cell type annotated assigned to each niche. Niche Table S5 was calculated based on the average number of neighbors and their cell type annotations for each niche. Niche type diversity was calculated using ‘scipy.stats.entrpopy’ and any niches that were more than 99% dominated by a single sample were excluded from analysis. T/NK and myeloid expression heatmaps were generated by averaging expression of cell-specific markers within all T/NK or myeloid cells from each niche. Any niches that contained less than 50 T/NK or 30 myeloid cells were excluded from heatmaps. Average niche expression was then zscored across all niches.

#### Niche marker abundance

To test differential expression of OX40 and CD57 in pre- and post-enriched niches, the top 5 enriched pre/post niches were compared. All T cells from each niche were aggregated and then post-DLI cell expression was compared to pre-DLI cell expression using a Mann-Whitney U test. For comparing responder pre-DLI expression of markers to nonresponder pre-DLI expression, all samples with more than half (>55%) pre-DLI cells were included from each response group, and a Mann-Whitney U test was used to compare marker expression between the two groups.

#### Niche proportion abundance

Proportion of cell types pre-DLI was calculated for all niches with >55% pre-DLI cells for both responder and nonresponder niches. T/NK, B, and myeloid cells were all tested separately using a Mann-Whitney U test between responder and nonresponder pre-enriched niches.

#### CODEX Image visualization

Marker visualization for individual windows was performed using QuPath^70^ imaging software.

### TCR analysis

T cells from BMMC, DLI products and one single patient peripheral blood mononuclear cells (PBMC, R3501) post-DLI were aggregated in one scanpy data object. Cells with a confident TCR beta chain were aggregated into clonotypes and clonality of each cell was calculated as the total number of times the CDR3 was observed across all sample time points and acquisition sites. Dimensionality reduction was performed by selecting highly variable genes using default analysis functions from scanpy except selecting only the first 15 principal components. Cells from two DLI products seemed to show strong signals related to heat shock events occasionally observed in thawing of cryopreserved cell suspensions. These differentially upregulated genes were identified using rank_gene_groups function from scanpy and removed from the highly variable genes list prior to dimensionality reduction. No other batch correction was performed on this subset of cells. To find genes correlated with clonality, Pearson correlation coefficient between vector of counts for each gene and vector of clonality across all cells were calculated. After Benjamini-Hochberg (BH) correction, significant correlations were selected (p value <.01 and correlation >.05 or <-.05). Differentially expressed genes between clusters 0 and 5 were calculated using rank_gene_groups function and a Wilcoxon Rank Sum test.

#### TCR Diversity

TCR diversity was calculated by measuring the GIni coefficient for all clones detected at each time point on a per-patient basis. Singletons were included in analysis. Significance was calculated using a Mann-Whitney U test (**Figure S6J**).

#### TCR VDJdb Analysis

VDJdb^85,86,166^ (v. 2023-04-26) was used, and analysis was limited to clones with clone size ≥2. Of 1,049 unique expanding clones, 36 clones overlapped with those in the VDJdb database (**Figure S6C**).

#### Tumor Specificity Score

Tumor specificity scores^87,88^ were calculated by taking the log 10 of total counts of all genes in each signature divided by the number of counts in each cell. CD8+ Tumor Specificity score calculated using following genes: *KRT86, RDH10, TYMS, HMOX1, GNG4, CXCL13, AFAP1L2, ACP5, MYO1E, LAYN, TNS3, TNFSF4, AKAP5, HAVCR2, ENTPD1, SLC2A8, ZBED2, MCM5, CAV1, GOLIM4, VCAM1, PON2, MTSS1, CD38, MS4A6A, TOX2, CSF1, GALNT2, FXYD2, PLPP1, LMCD1, MYL6B, LAG3, HLA-DRA, IGFLR1, CCDC50, CD27, KIAA1324, CDKN2A, CD70, ABHD6, CTLA4, PDCD1, GEM, NUSAP1, TOX, CXCR6, NMB, HOPX, CLIC3, INPP5F, SNAP47, TSHZ2, HLA-DMA, SIT1, HLA-DRB1, TUBB, PYCARD, ADGRG1, HLA-DQA1, PRF1, HLA-DPA1, PTMS, CKS1B, HIPK2, CHST12, LSP1, FAM3C, SLC1A4, NUDT1, DNPH1*. CD4+ Tumor Specificity score calculated using following genes: *ADGRG1, CADM1, RDH10, HMOX1, GNLY, HAVCR2, PRF1, SLC27A2, NKG7, CCL4, GZMA, SLC1A4, GZMK, LAG3, NCDN, CST7, CXCR6, UBE2E3, KIAA0319L, CHST12, PDCD1, SLA2, PTMS, CCL5, RHBDF2, CD27, IGFLR1, CTSW, GGA2, APOBEC3G*

#### TCR post-DLI enrichment analysis

TCR data was limited to clones existing in post-DLI samples with total clone size ≥2 across all time points. Overlap was calculated per subject based on the number of unique clones present in post-DLI that were also present pre-DLI or in the DLI product (**Figures 6F** and **S6G**-**I**). Fisher’s exact test (FET) was performed on each patient by designing a 2×2 contingency table with columns 1 and 2 representing clonotype existing in post or not existing in post sample, respectively, and rows 1 and 2 reflecting clones detected at pre-DLI, and in the DLI product. FET was calculated on each patient separately.

### External validation of *ZNF683-*expression T cells in AML bone marrow

Previously published scRNA-seq data analyzed here was found at Gene Expression Omnibus with the accession code GSE198052. Data was read into Seurat (version 4.1.3)^167^ and filtered for number of features between 200 and 2500 as well as under 5% mitochondrial RNA expression. The data was then normalized, scaled, and processed using the NormalizeData, FindVariableFeatures, ScaleData, and RunPCA functions in Seurat. Data was filtered into two Seurat objects: one comprising only T cells (consisting of already assigned labels ‘CD4’, ‘CD8’, and ‘Unconventional T’), and one comprising T cells and NK cells (consisting of the aforementioned labels as well as ‘NK’ and ‘preT/NK’). Healthy and stable disease patient samples were filtered out to avoid bias. Each object was separately normalized and scaled as previously mentioned and then clustered to create UMAPs using the FindNeighbors, FindClusters, and RunUMAP functions in Seurat. Heat maps and dot plots were generated showing average gene expression (z-score) for a given cluster. In the cases comparing responder and non-responder, the data was separated and then averaged to show gene expression. Frequency graphs were calculated using the number of cells per cluster in each patient.

### External validation Transcriptome comparison

Previously published scRNA-seq data^55,71,72,75^ were obtained and subsetted to contain cells of interest (CD8 T cells and NK cells if available). Meta data including Normal/Malignant or HSCT status were merged with cluster names. A celltypist^142^ logistic regression model was trained on each dataset based on these cluster names. Each model was used as a reference to predict the most likely CD8 and NK cells from the BMMC subset of the AML cohort.

## Supplemental Figures

**Figure S1.**
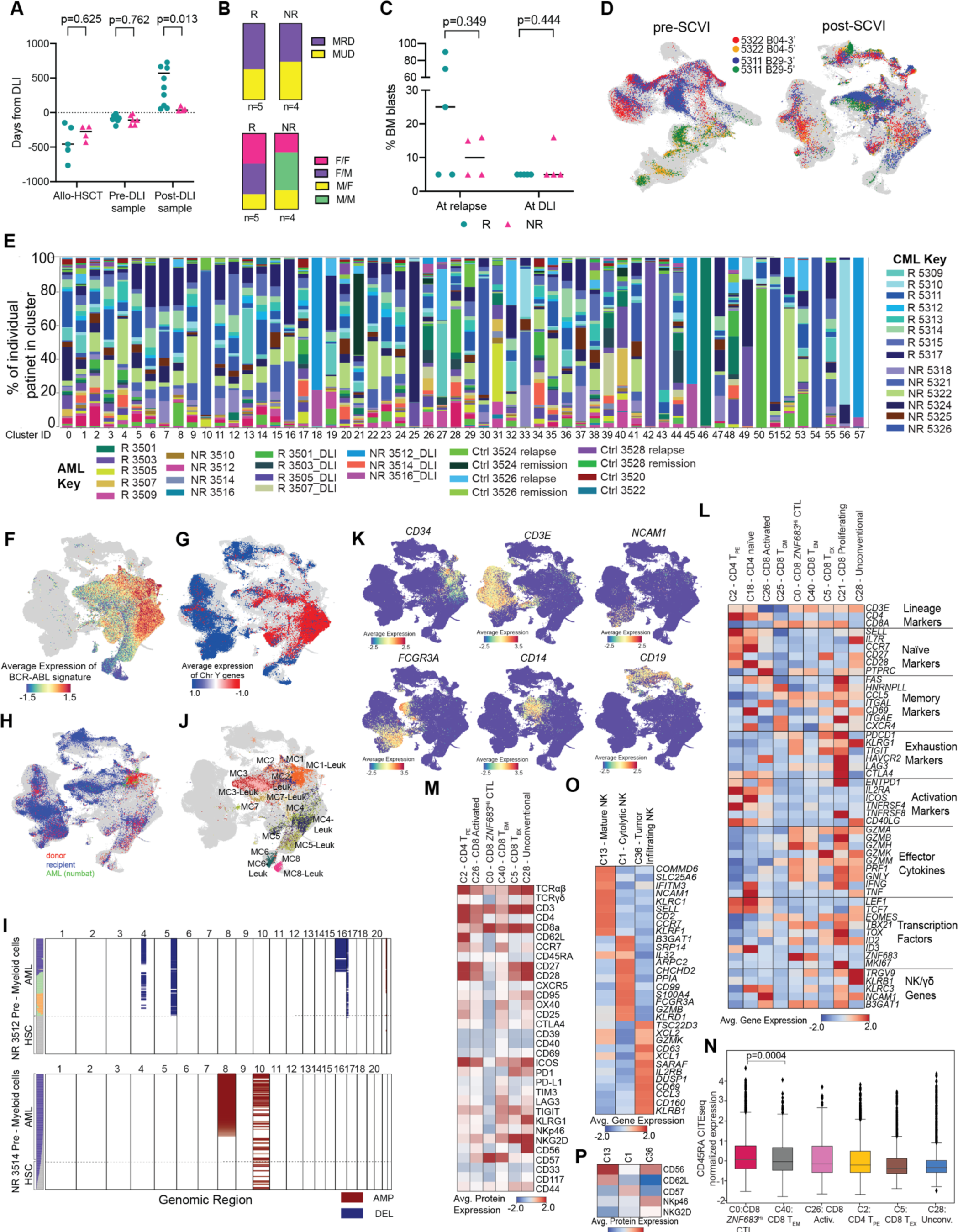
Details of clinical samples and global map. **A.** Time from HSCT to relapse for each patient and time of each AML bone marrow biopsy sample analyzed relative to DLI (middle dotted line). **B.** *Left*: Bar graphs of proportion of matched related donor (MRD) and matched unrelated donor (MUD) for responders (R, left bar) and nonresponders (NR, right bar). *Right*: Bar graphs of patient/donor sex for R (left bar) and NR (right bar). **C.** Leukemia burden, as measured by percent of bone marrow (BM) blasts, at the time of relapse (*left*) and at the time of DLI (*right*) for R (dark teal) and NR (pink). **D.** 2D UMAP projection of all merged cells before (*left*) and after (*right*) batch correction with scVI. Cells from samples profiled with both 3’ and 5’ scRNA-seq chemistries are shown in color and other cells are shown in gray. **E.** Percent of cells from each subject within each cluster. **F.** 2D UMAP coloring myeloid cells by expression of BCR-ABL translocation signature used to identify clusters enriched in CML leukemia cells. **G.** Coloring of 2D UMAP projection of all cells by average expression of chromosome Y genes in samples from patients of the opposite sex from their HSCT donor. **H.** Coloring of 2D UMAP with AML sample cells of donor (red) or recipient origin (blue) as determined by Vireo, with cells determined to be of leukemic origin by Numbat in green. **I.** Heatmap of CNVs predicted from scRNA-seq data of two NRs (3512, 3514) using Numbat. Rows indicate myeloid cells from pre-DLI samples, columns represent genomic regions. Amplification and deletion are shown in red and blue respectively. **I.** Coloring of 2D UMAP myeloid cells from AML subjects by metacluster. **K.** Coloring of 2D UMAP by expression of major cell type markers. **L.** Average expression of T cell lineage defining genes as well as other genes of interest to delineate T cell functional subset. Red: higher expression, blue: lower expression. **M.** Average protein expression by scCITE-seq for T cell clusters with >1000 cells having scCITE-seq data. **N.** Box plot of normalized CITE-seq expression of CD45RA among T cell clusters. (C0 vs C40 p=0.0004, Kolmogorov Smirnov test). **O.** Average expression of NK cluster lineage defining and functional genes. **P.** Average protein expression by scCITE-seq in the three NK cell clusters.

**Figure S2.**
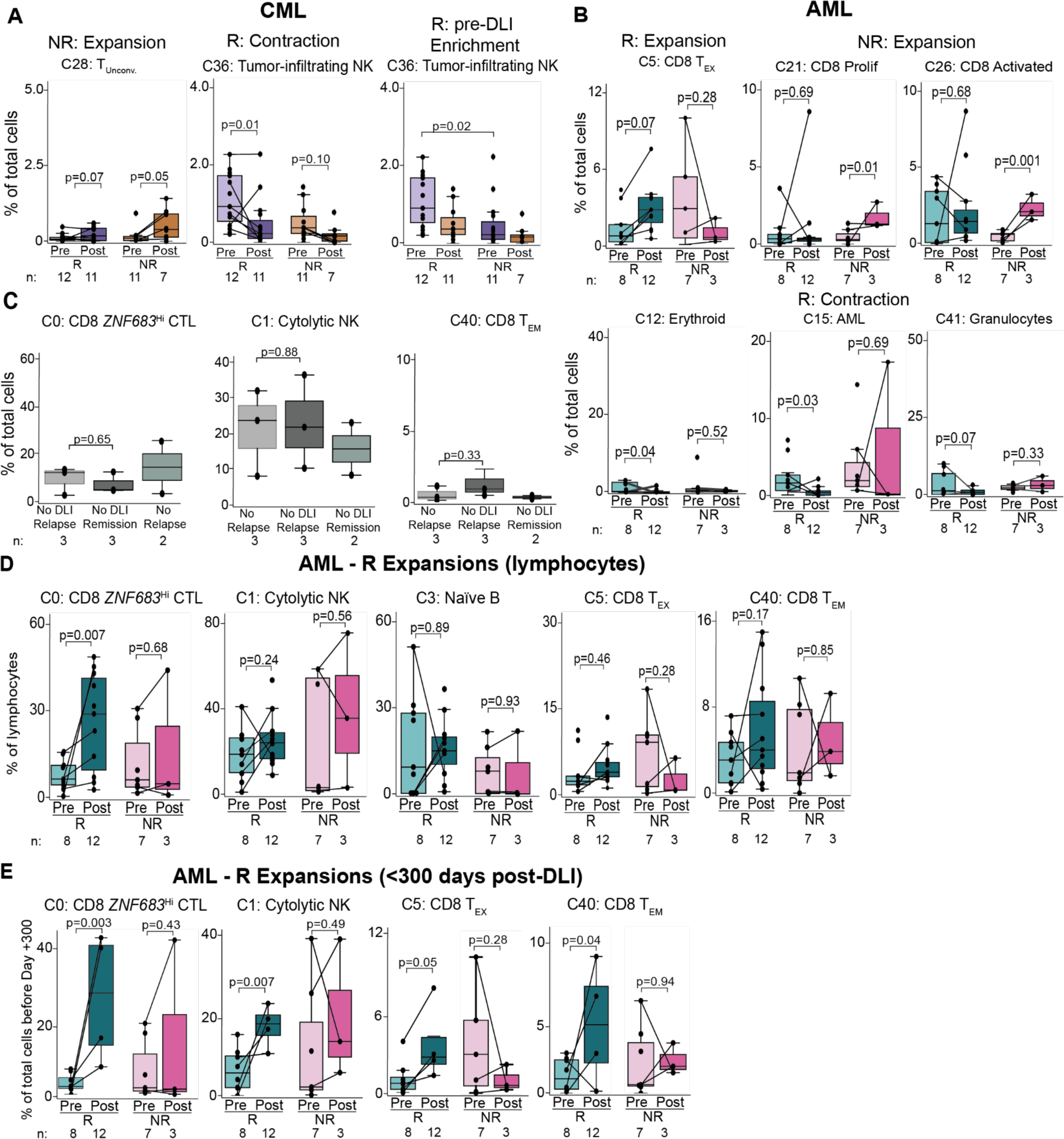
Immune cell states enriched between clinical groups. **A.** Percent of all cells for each CML cluster with significant expansion or contraction in R or NRs that were not shown in Figure 2A. **B.** Same as (A) for clusters with significant expansion or contraction in AML Rs or NRs. **C.** Percent C0 (left), C1 (middle) and C40 (right) cells of all cells from control patients. Paired relapse/pre-treatment and remission/post-treatment samples for chemotherapy only/no DLI controls. For no relapse controls, only one sample per patient was analyzed. **D.** Percent of all cells of total lymphocytes (excluding myeloid compartment and leukemia cells to control for disease burden) for clusters with significant expansion in AML. **E.** Same as (D) with limiting post-DLI samples to those collected within 300 days of DLI.

**Figure S3.**
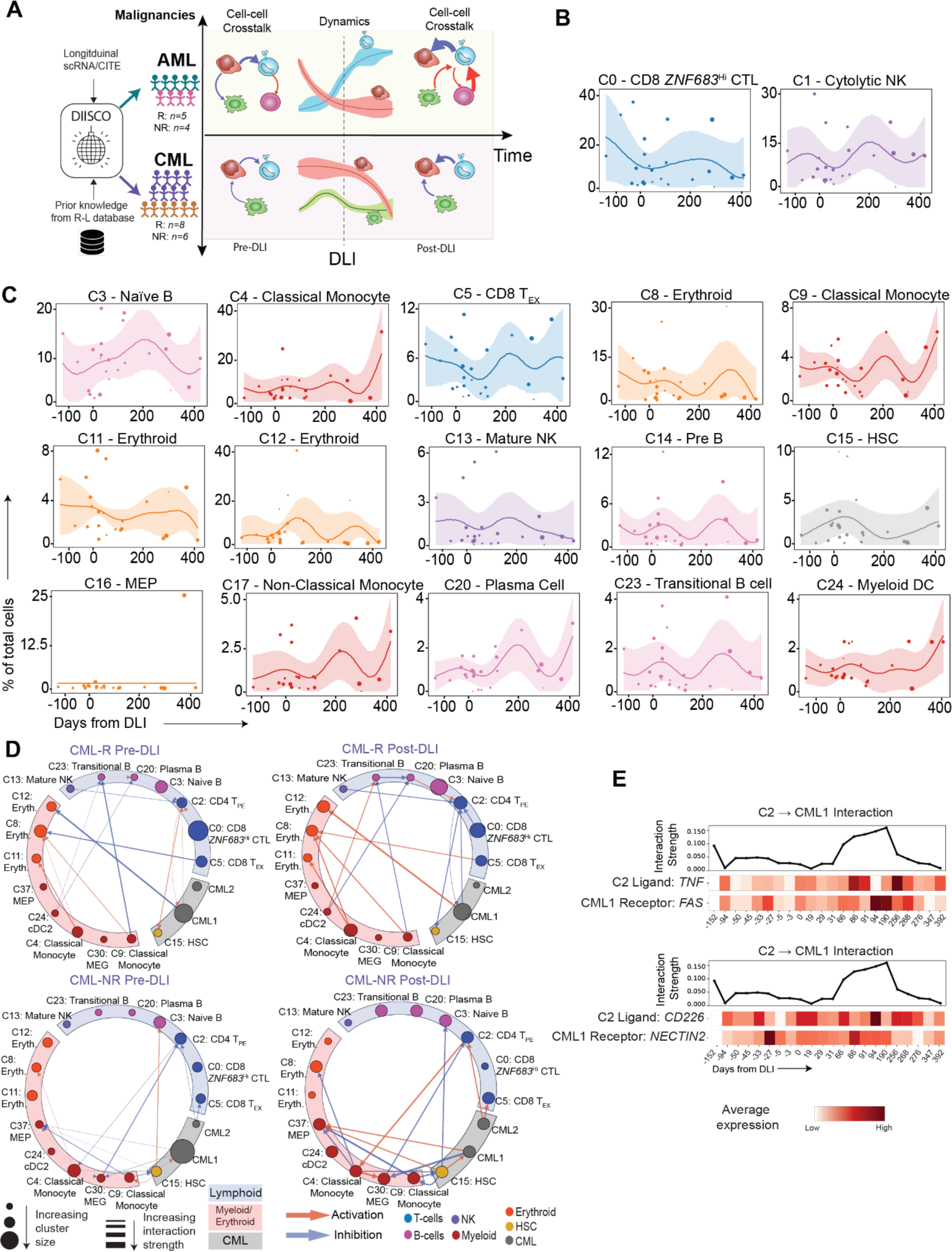
Details of predicted intercellular interactions in CML. **A.** Schema of DIISCO for AML and CML. **B.** DIISCO predictions for CML showing dynamics for clusters with significant dynamics in AML only. **C.** DIISCO predictions for remaining clusters in CML responders not shown in (B) or Figure 2B. **D.** DIISCO interactions predicted between C2: CD4 T_PE_, C13: Mature NK, and C15: HSCs for R, but C2-myeloid interactions in NR. Predicted activating interactions are indicated by red arrows while inhibitory interactions are indicated by blue arrows. **E.** Examples of the top predicted interactions betwee_n C_2: CD4 T_PE_ and CML1.

**Figure S4.**
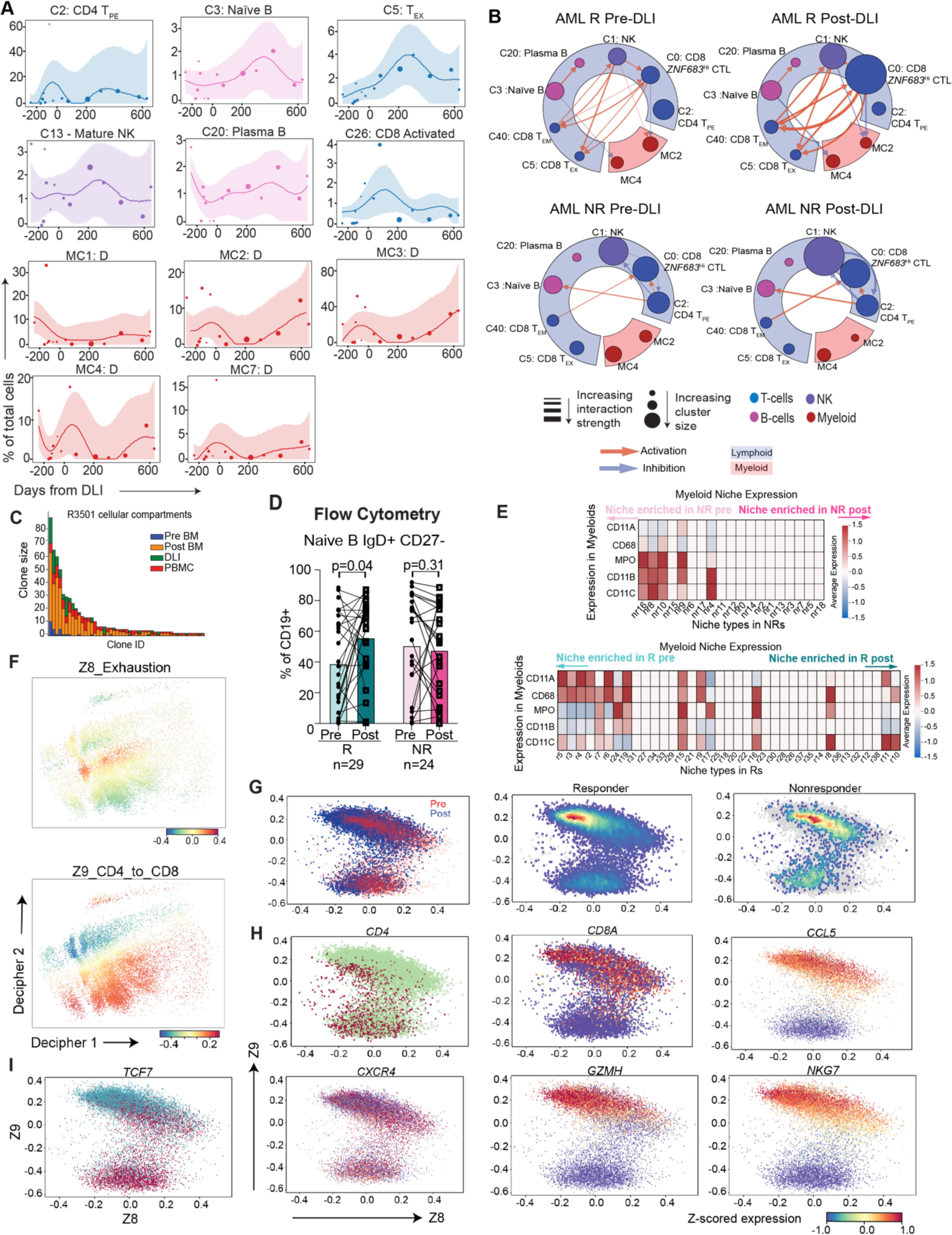
C0 CD8 *ZNF683*^Hi^ CTLs are a central hub for immune activity predicted by DIISCO. **A** DIISCO predictions for the remaining clusters not shown in **Figure 2D** for R. *Top*: From left to right, C2 (CD4 T_PE_), C3 (Naive B), C5 (CD8 T_EX_), C13 (Mature NK), C20 (Plasma B cell), C26 (CD8 Activated). *Bottom*: Metacluster 1 (MC1, donor cells), metacluster 2 (MC2, donor cells), metacluster 3 (MC3, donor cells), metacluster 4 (MC4, donor cells), metacluster 7 (MC7, donor cells). Days from DLI on X-axis, % of total cells in each sample on Y-axis. **B.** DIISCO prediction network limited to only immune cells (all patient derived myeloid/leukemia cells excluded). **C.** Top expanded T cell clones for R3501 in different compartments (i.e. BM [pre-DLI: blue, post-DLI: orange], DLI product [green], and peripheral blood mononuclear cells [PBMC, red] at the same time point as the post-DLI BM). Each bar represents one clone. Y axis indicates the number of cells found in that clone across time. **D.** Percent of naive B cells by flow cytometry of PBMC samples from an independent cohort of post-HSCT relapsed AML patients who received DLI including 29 R and 24 NR. **E.** Expression of CODEX markers for myeloid cells in each niche identified in Rs (*left*) and NRs (*right*). **F.** 2D visualization of Decipher components (Decipher 1 and Decipher 2) as x,y coordinates, colored by components Z8 (exhaustion) and Z9 (CD4 vs CD8). **G.** T cells plotted with Z8 and Z9 as x,y coordinates. *Left*: Cells colored by pre (red) and post (blue). *Middle*: R cells colored by density - red indicates higher density. *Right*: NR cells colored by density - red indicates higher density. **H-I.** T cells plotted with Z8 (x axis) and Z9 (y axis) components colored by gene expression of *CD4, CD8A, CCL5, CXCR4, GZMH, NKG7* (**H**) and *TCF7* (**I**). Red indicates higher expression.

**Figure S5.**
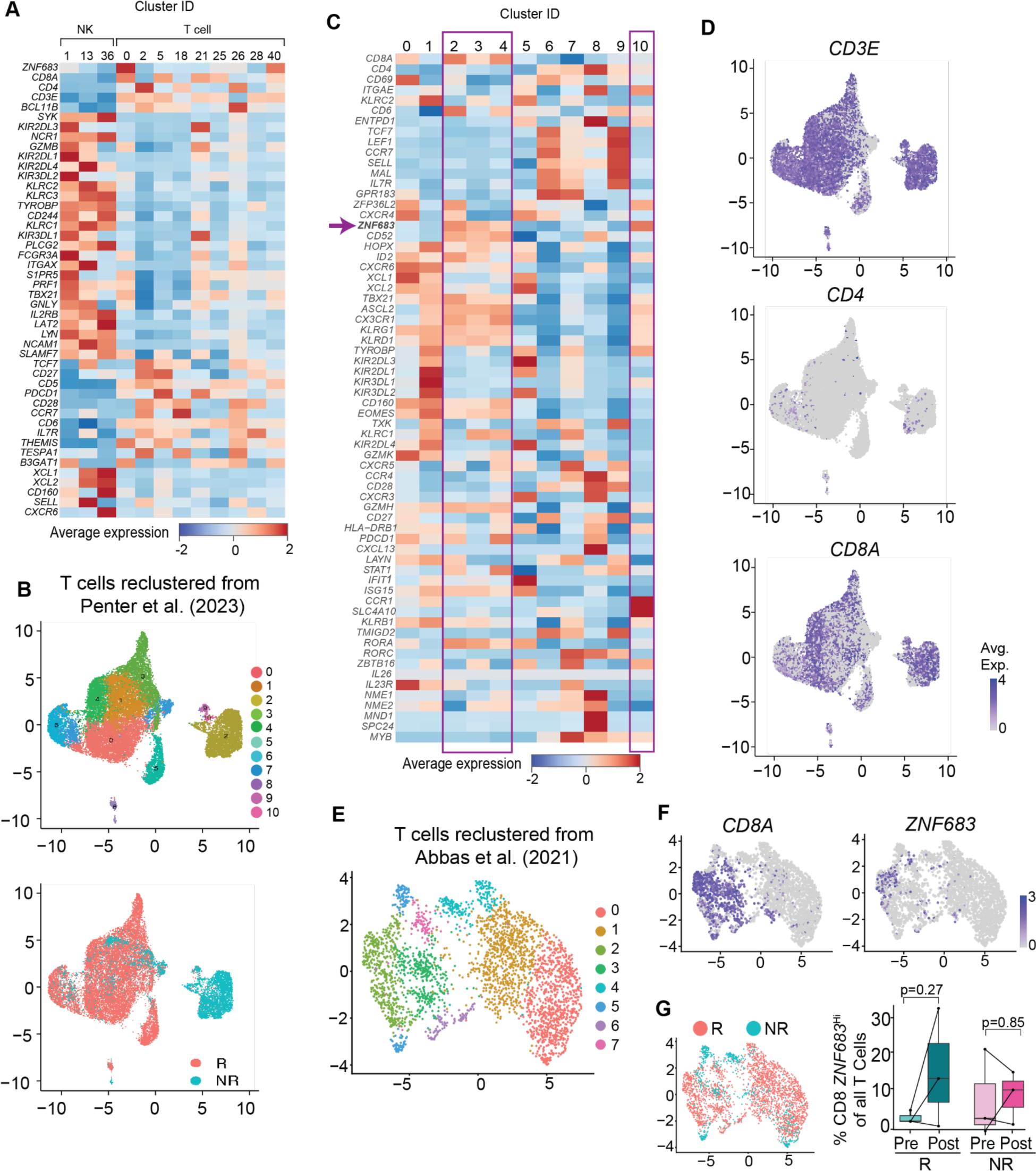
Comparison of T and NK cell clusters to external datasets. **A.** Comparison of all NK and T cell clusters to a previously reported signature for NK-like CD8+ T cells (Sottile et al, *Sci Immunol*, 2021). **B.** 2D UMAP with reclustering of the T cells from Penter et al, *Blood*. 2023. *Top:* Eleven distinct clusters were identified. *Bottom:* Coloring of 2D UMAP by response to CTLA-4 blockade. **C.** Average expression of T cell lineage-defining genes and markers for phenotypic/functional characterization from (B). **D.** 2D UMAPs of T cells from (B) showing expression of *CD3E* (*top*), *CD4* (*middle*), *CD8A* (*bottom*). **E.** 2D UMAP with reclustering of T cells from Abbas et al, *Nat Commun*, 2021. **F.** 2D UMAP of cells from (E) showing average expression of *CD8A* (*left*) and *ZNF683* (*right*). **G.** *Left:* 2D UMAP of (**E**) showing colored by response (R, salmon pink) or nonresponse (NR, blue) to PD-1 blockade. *Right:* Percent of CD8+ *ZNF683*^Hi^ cells of total T cells pre-treatment (lighter colors) and post-treatment (darker colors) for R (teal, p=0.27, paired t test) and NR (pink, p=0.85, paired t test).

**Figure S6.**
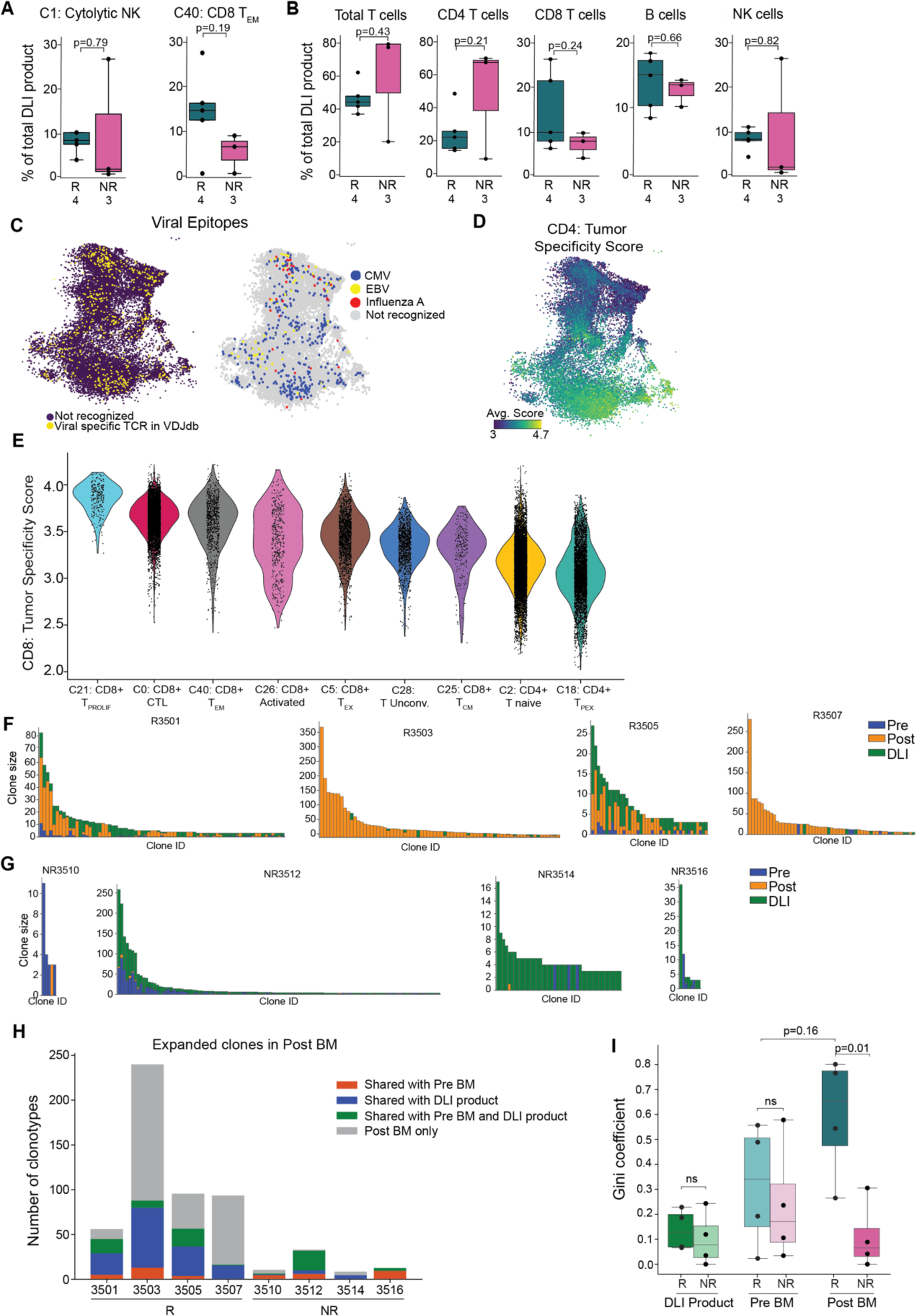
T cell clonality and specificity. **A.** Percent of C1 and C40 of all DLI product cells per patient for R (teal) and NR (pink). **B.** Same as (A) comparing percent of major cell types in DLI product cells in R and NR. **C.** Specificity of TCR clones compared to known viral epitopes identified by VDJdb. *Left:* Yellow dots - TCR specific to a known viral epitope. *Right:* Clones specific for known cytomegalovirus (CMV, blue), Epstein-Barr virus (EBV, yellow), or Influenza A (red) viral antigens. **D.** Coloring of T cells by similarity to a previously published CD4 tumor specificity score. Yellow - higher specificity score; blue - lower specificity score. **E.** Violin plot of each T cell cluster ordered by most to least similar to the CD8 tumor specificity score **F.** Top expanded T cell clones for each individual R. Each bar represents one clone. Y axis indicates the number of cells found in that clone across time (pre-DLI [blue] and post-DLI [orange]) and compartments (DLI product [green] versus bone marrow). **G.** Top expanded T cell clones for each individual NR. Axes and colors are the same as in (F). **H.** Number of expanded clonotypes shared between DLI product and post-DLI BM (blue) versus those shared between pre-DLI BM and post-DLI BM (orange), clones found in all three sample types (green) as well as expanded clones in the post BM that were not found in pre BM or DLI product (grey) for each subject. **I.** Gini coefficient for T clonotype diversity among DLI products of R(dark green) vs NR (light green, p=0.59), pre-DLI BM for R (light teal) vs NR (light pink, p=0.6), and post-DLI BM for R (dark teal) vs NR (dark pink, p=0.01) (t test).

**Figure S7.**
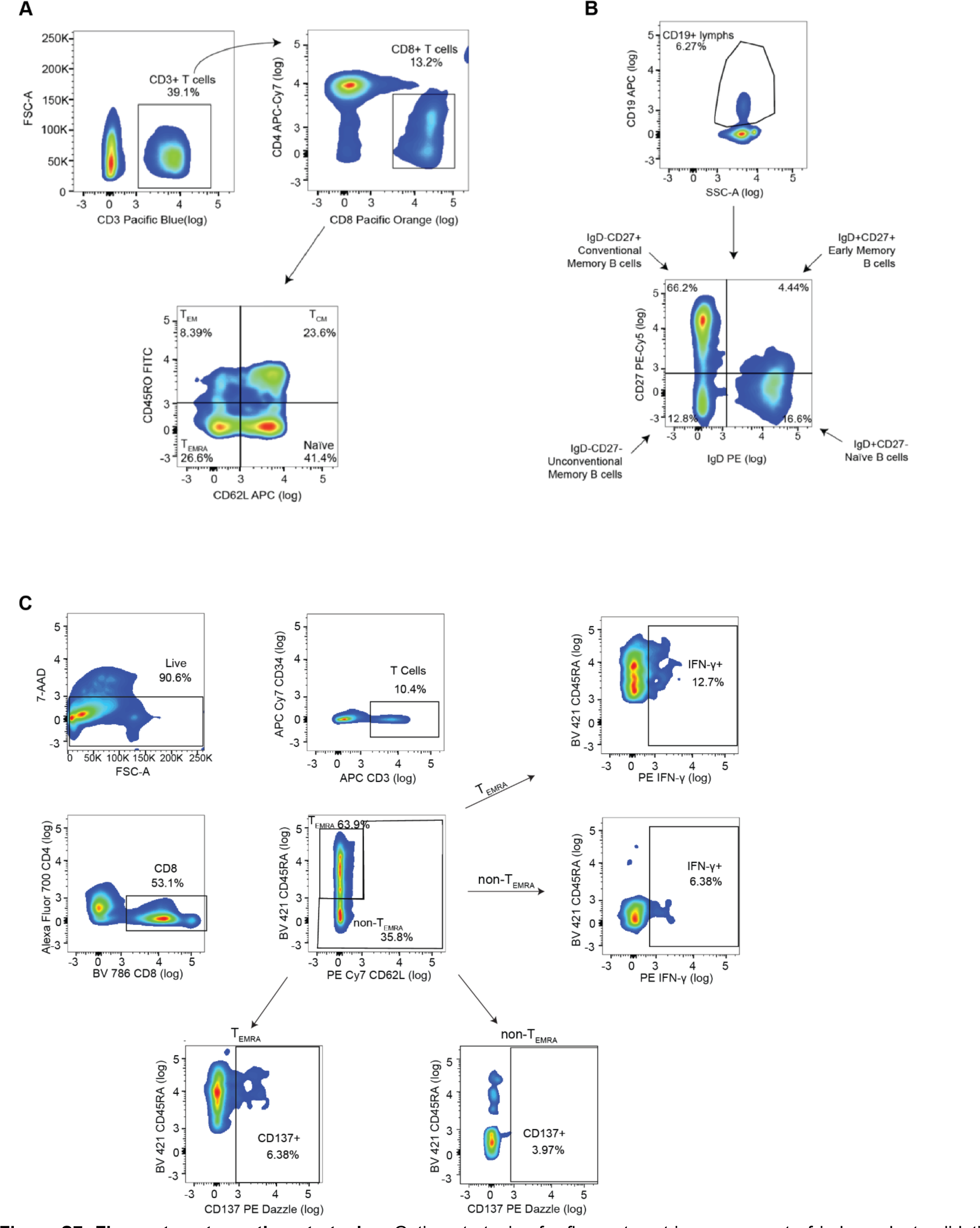
Flow cytometry gating strategies. Gating strategies for flow cytometric assessment of independent validation cohort peripheral blood CD8 T_EMRA_ CTLs (A) and naive B cells (B). C. Gating strategy for *in vitro* CD8 T_EMRA_ CTL activation assay. CD8+ T cells were gated into T_EMRA_ (CD45RO-CD62L-) or non-T_EMRA_ (all other CD8 T cells) subsets. Activation was measured by expression of CD137 or IFN-ɣ (Miltenyi) in separate experiments.

